# The Lipid Hydrolase ABHD6 is a Therapeutic Target in Metabolic Dysfunction-Associated Steatotic Liver Disease (MASLD)-Related Hepatocellular Carcinoma

**DOI:** 10.1101/2025.07.29.667509

**Authors:** Danny Orabi, William J. Massey, Kevin K. Fung, Venkateshwari Varadharajan, Iyappan Ramachandiran, Rakhee Banerjee, Daniel J. Silver, Lucas J. Osborn, Amanda L. Brown, Stephanie Marshall, Daniel Ferguson, Shijie Cao, Rebecca C. Schugar, Chelsea Finney, Chase Neumann, Amy C. Burrows, Anthony J. Horak, Preeti Pathak, Robert N. Helsley, Dominik Bulfon, Robert Zimmermann, Yat Hei Leung, S.R. Murthy Madiraju, Marc Prentki, Richard G. Lee, Adam E. Mullick, Olga Zergeeva, Srinivasan Dasarathy, Zhenghong Lee, Daniela S. Allende, Federico Aucejo, Justin D. Lathia, J. Mark Brown

**Author notes:** <u>To whom correspondence should be addressed:</u> Danny Orabi, M.D. Ph.D.; or J. Mark Brown, Ph.D.

## Abstract

Primary liver cancer accounts for approximately 700,000 deaths worldwide annually ranking third in cancer-related mortality, with hepatocellular carcinoma (HCC) comprising the majority of these tumors. Metabolic dysfunction-associated steatotic liver disease (MASLD) is currently a leading cause of HCC in the United States. We previously identified the lipid hydrolase alpha/beta hydrolase domain 6 (ABHD6) as a key mediator of the development of metabolic syndrome and intimately involved in cell signaling, making it a prime target for investigation in MASLD-related HCC. ABHD6 displays higher expression within HCC tumor cores when compared to adjacent non-tumor liver tissue in human subjects. Using an in vivo antisense oligonucleotide (ASO)-driven knockdown approach, we have shown the inhibition of ABHD6 prevents the development and progression of HCC in an obesity/MASLD-driven mouse model. Additionally, a xenograft model using the human Huh7 cell line displayed reduced tumor engraftment and growth with ABHD6 genetic deletion and small molecule inhibition. ABHD6 knockout cells demonstrated increased levels of bis(monoacylglycerol)phosphates (BMPs), lipids relevant to high fat diet-induced lysosomal dysfunction, and knockout cells also demonstrated altered autophagy and lysosomal activity using *in vitro* model of saturated fatty acid-induced lipotoxicity. These studies reveal novel lipid signaling mechanisms by which MASLD progresses towards HCC and provide support for ABHD6 as a therapeutic target in HCC.

**Significance:** We have identified that alpha/beta hydrolase domain 6 (ABHD6) plays a role in lysosomal membrane lipid remodeling pathways that are relevant in obesity/MASLD-driven HCC. Inhibitors targeting ABHD6 reorganize lysosomal lipid homeostasis to improve outcomes in HCC.

## Introduction

Primary liver cancer currently sits as the third leading cause of cancer-related mortality worldwide, with hepatocellular carcinoma (HCC) comprising the great majority of these tumors [1]. The five-year survival for HCC remains under 20% and there is a dire need for advancements in treatment [2]. HCC typically arises in a background of chronic liver disease most often from chronic viral hepatitis infection, alcohol, and metabolic dysfunction-associated steatotic liver disease (MASLD) [3]. The epidemiologic landscape of liver disease and HCC has seen large shifts over the past few decades with MASLD quickly moving to the forefront [3–6]. The liver plays a key role in lipid processing with several intricate pathways working in tandem to achieve a delicate metabolic homeostasis. Liver steatosis and MASLD progression results from excess dietary lipid intake, maladaptive adipose tissue signaling, insulin resistance, and aberrant hepatocyte lipid trafficking [7]. The resultant hepatic and systemic metabolic dysfunction promote HCC through chronic inflammation, DNA mutations, gene expression changes, cell cycle modulation, and immune dysregulation [8,9]. Treatments targeting lipid handling in MASLD and HCC may provide novel avenues of treatment, and finding a mechanism common to both would serve in both treatment and chemoprevention [10].

The serine hydrolase alpha/beta hydrolase domain 6 (ABHD6) was initially found to play a role in endocannabinoid (ECB) signaling, given that ABHD6 can hydrolyze the key ECB 2-archidonoylglycerol (2-AG) [11]. Shortly thereafter, its various roles in metabolic homeostasis and cancer were established. It was found to play a key role in high fat diet-induced metabolic syndrome and liver steatosis, pancreatic beta cell insulin release, and adipose tissue physiology and inflammation [12–18]. A study that screened over 700 drugs for inhibition of pancreatic cancer metastasis found two ABHD6 inhibitors to be the leading candidates [19]. A subsequent study found ABHD6 inhibition reduced non-small cell lung cancer metastasis in a mouse model via monoacylglycerol (MAG)-mediated reduction in epithelial-to-mesenchymal transition [20]. Substrates of ABHD6 include MAG and bis(monoacylglycerol)phosphate (BMP) lipids [13,21,22]. While monoacylglycerol lipase (MAGL) is highly expressed in the liver and likely accounts for the majority of hepatic MAG hydrolysis, ABHD6 is the principle enzyme responsible for hepatic BMP metabolism [22,23]. BMP lipids localize to the late endosome/lysosome [24]. Hepatic and plasma BMP lipids increase in response to lysosomal storage disorders and high fat diet feeding [23]. The interrelationship between ABHD6, BMP lipids, MASLD, and lysosomal pathways influenced this study’s line of investigation. Lysosomal pathways, notably autophagy and the endolysosomal network, play intimate roles in lipid metabolism, cell signaling, and cell cycle regulation and have been linked to both MASLD and HCC making it a promising therapeutic target [25–33]. In this study, we explore the role of ABHD6 in MASLD-related HCC. We used two separate mouse models to demonstrate that inhibition of ABHD6 reduces tumor development and progression. ABHD6 inhibition led to an increase in cellular BMP lipid species, and to altered autophagy and lysosomal activity in a cell culture model of saturated fatty acid-induced lipotoxicity.

## Materials and Methods

### Human HCC Tissue and ABHD6 Immunohistochemistry

A repository of paraffin embedded liver specimens from patients undergoing resection of HCC at the Cleveland Clinic Foundation were analyzed for ABHD6. A maltose binding protein ABHD6 fusion protein construct was generated to create affinity-purified rabbit polyclonal antibodies against murine ABHD6 as previously described [12]. ABHD6-directed immunohistochemistry (IHC) was performed using this rabbit ABHD6 antibody diluted 1:85 using SignalStain Diluent (Cell Signaling Technology) and incubated for 48 minutes, ambient. Automated staining used the VENTANA Discovery XT platform, Res IHC Omni-UltraMap HRP XT software, and was detected using OMap anti-Rb HRP (Ventana Medical Systems Inc. Tucson, AZ). Antigen Retrieval selected standard Cell Conditioning#1. VENTANA’S DISCOVERY Antibody Block (RUO) incubation lasted 20 minutes, and the multimer HRP was selected for 20 minutes. The negative tissue control was an ABHD6 knock-out mouse liver block obtained from our laboratory. Positive tissue controls included gastric mucosa, brain tissue, and liver. Histological analysis and patient chart review were performed by a board-certified pathologist with expertise in hepatobiliary pathology and a resident physician in the Department of Pathology at the Cleveland Clinic Foundation. ABHD6-directed IHC slides were scored by intensity of staining (whether visible at 4x, 10x, or 20x magnification, or not visible), and percent of tumor area staining (<5%, 5-50%, or >50% tumor staining).

### Animal Studies

C57BL/6J and Rag2 knockout (C57BL/6N) mice were purchased from Jackson Laboratories (Bar Harbor, ME USA) and Taconic Biosciences (Rensselaer, NY USA), respectively. Mice were fed standard rodent chow diet ad libitum unless otherwise specified in the experimental outlines described below. High fat diet (HFD) had a 60% kCal from fat composition (diet D12492 from Research Diets Inc). Mice were housed in the Biological Resources Unit of the Lerner Research Institute, Cleveland, OH USA. Mice were maintained in an Association for the Assessment and Accreditation of Laboratory Animal Care, International-approved animal facility, and all experimental protocols were approved by the Institutional Animal Care and use Committee of the Cleveland Clinic.

### Obesity/MASLD-Driven Mouse Model of HCC

C57BL/6J mice (strain# 000664) were purchased from Jackson Laboratories (Bar Harbor, ME USA). Males and females were paired to produce timed litters. As described by Yoshimoto et al. [34], 4-5-day old pups were given a single treatment of a 50 μl solution of 0.5% DMBA (7,12-dimethylbenz(a)anthracene, Sigma) in acetone painted onto their dorsal surface with a P200 pipette tip [34]. At 21 days of age, male pups were weaned to separate cages and started on HFD. Of note, only male mice were studied as females did not develop liver tumors using this model in our experience (data not shown), consistent with what has been previously reported [34,35]. In the HCC development experiment, upon weaning, mice were treated bi-weekly with 25 mg/kg of an ABHD6 targeting antisense oligonucleotide (ASO) or a scrambled non-targeting control ASO (sequences available on request) as previously described [12]. After 30 weeks on HFD i.e. 33 weeks of age, mice were euthanized. Images of ventral and dorsal surfaces of all liver lobes were taken, and image analysis performed by two independent data reviewers. Lung and liver tissue were harvested as flash frozen specimens and formalin fixed specimens. The formalin fixed specimens were then paraffin-embedded and Hematoxylin and eosin (H&E) stained as previously described [12], then scored for tumor burden by a board-certified pathologist. In the HCC progression experiment, pups were treated with DMBA then weaned to HFD as described above. At 25 weeks of age, mice underwent weekly ultrasound imaging to assess tumor burden. Mice were matched by tumor burden then randomized to receive bi-weekly injections of either an ABHD6-targeting ASO or a scrambled non-targeting control ASO. Tumor locations were documented and weekly ultrasound was performed to monitor specific tumor lesion progression until endpoint, defined by poor physiologic status or inability to measure tumor nodules. Mice were then euthanized and liver tumor and non-tumor tissues flash frozen for later analysis.

### Orthotopic Xenograft Model of HCC

C57BL/6N Rag2 knockout (model RAGN12) male and female mice were purchased from Taconic Biosciences (Rensselaer, NY USA) and bred in house. At 3 weeks of age, male mice were weaned into cages of no more than 4 mice per cage. Female mice were not used given their markedly low rates of tumor engraftment (data not shown). At 15-20 weeks of age, mice were started on HFD and maintained on HFD throughout the remainder of the experiment i.e. until time of necropsy to help recapitulate a MASLD phenotype. After 2 weeks of HFD feeding, an orthotopic xenograft was performed via direct liver intraparenchymal injection. Surgery was performed using a Leica Wild M650 surgical microscope (Wetzlar, Germany) under 6x – 25x magnification. Briefly, induction and maintenance of anesthesia were achieved with 2-4% and 1-3% isoflurane, respectively. Antibiotic-free eye ointment was applied. The abdomen was shaved and cleaned with betadine and 70% alcohol. Bupivacaine (2 mg/kg) was used for local anesthesia and a midline laparotomy was performed. Using a 31-gauge insulin syringe needle, 1 million luciferase-transduced cells in single cell suspension in 50 μL of serum-free antibiotic-free Dulbecco’s Modified Eagle Medium (DMEM) culture media (Media Core, Lerner Research Institute, Cleveland, OH) were injected into the inferior aspect of the left liver lobe with care to prevent leakage from the liver capsule. This was done by skiving the needle superficially just inferior to the liver capsule for 3 mm, then penetrating slightly deeper and slowly injecting cells over 1 minute. Failure of the liver capsule to fracture and blanch was considered evidence for vascular communication in which case engraftment was unlikely and the mouse was excluded from the experiment and euthanized. After removal of the needle, a cotton swab applicator was gently placed over the injection site for 2 minutes. Another intra-parenchymal injection of 1 million luciferase-transduced cells was performed in the inferior aspect of the right side of the median liver lobe using the same technique. The abdominal wall was closed with 6-0 PDS absorbable suture in a simple continuous fashion, and the skin was closed with 5-0 polypropylene non-absorbable suture in a horizontal-mattress interrupted fashion. Mice were recovered and maintained on HFD for the remainder of the experiment. Postoperative analgesia was maintained with an intraoperative dose of buprenorphine (0.1 mg/kg) and meloxicam (2 mg/kg), followed by twice-daily administration of buprenorphine (0.1 mg/kg) for two days postoperatively. Polypropylene skin stitches were removed approximately 10 days after surgery.

In the tumor engraftment experiment, mice in each cage were randomized to receive either wild type (WT) control or ABHD6Δ Huh7 cells. Bioluminescence imaging was performed at 2, 3.5, and 5 weeks postoperatively. Ultrasound was performed once at 5 weeks postoperatively to assess tumor size. At 5 weeks postoperatively, mice were euthanized and tumor was harvested as flash frozen specimens and formalin fixed specimens. Note: in the rare event of extrahepatic tumor formation (one event total in this study), the mouse was excluded from data analysis. In the tumor progression experiment, the surgical procedure was slightly modified such that a 2 mm x 2 mm piece of Surgicel Nu-Knit hemostatic gauze (item no. 1946) was left over the needle puncture site to reduce cell leakage and increase rate of engraftment. In this experiment, only WT Huh7 cells were used. One week after surgery, mice in each cage were randomized to receive 1 mg/kg KT203 (Cayman Chemical, item no. 14819), a peripheral (without penetration into the central nervous system) irreversible small molecule inhibitor of both mouse and human ABHD6 [36], or vehicle alone (18:1:1 saline:etoh:Kolliphor/PEG40 (Sigma, item no. 07076)) via daily intraperitoneal injection. Bioluminescence imaging was performed at 2, 3, and 4 weeks as described above. Ultrasound imaging was performed at 3 weeks, 3 weeks and 5 days, and 4 weeks and 3 days (five-day intervals). Mice were then euthanized and tumor was harvested as flash frozen specimens and formalin fixed specimens.

### Ultrasound Imaging

Induction and maintenance of anesthesia was achieved with 2-4% and 1-3% isoflurane, respectively. Mice were placed on a warmed stage and imaged using either an ACUSON S2000, HELX Evolution (Siemens) ultrasound with a 12L4 transducer or a Vevo 2100 (Visual Sonics) ultrasound with an MS550S transducer (supported by NIH Project 1S10OD021561-01A1) in the DMBA and xenograft mouse models, respectively. Tumors were imaged in the frontal and sagittal planes to generate a mediolateral diameter, craniocaudal diameter, and two anteroposterior diameters. Mice were recovered and returned to their cage mates. Tumor volume was calculated using the equation 4/3 × π × (mediolateral radius) × (craniocaudal radius) × (average anteroposterior radius).

### Bioluminescence Imaging

Huh7 cells were stably luciferase transduced as described below. *In vivo* grade luciferin (VivoGlo, Promega, item no. P1043) was purchased and suspended in 0.9% normal saline as a 15 mg/mL solution, aliquoted, and stored at -20 degrees. Induction and maintenance of anesthesia was achieved with 2-4% and 1-3% isoflurane, respectively. An intraperitoneal injection of 75 mg/kg luciferin suspended in 150 uL of 0.9% normal saline was administered. Five minutes after injection, mice were placed on a warmed stage and imaged using an IVIS Spectrum In Vivo Imaging System (PerkinElmer, Waltham, Mass) every 2 minutes for 10 minutes total. Mice were recovered and returned to their cages.

### Cell Culture, Lentiviral Transduction, and CRISPR-Cas9 Genome Editing

The human HCC cell line Huh7 obtained from the Lerner Research Institute Cell Services Core, Cleveland, OH were confirmed to be mycoplasma negative, and cell authentication via short tandem repeat analysis was performed by IDEXX BioAnalytics (North Grafton, MA). Cells were cultured in Dulbecco’s Modified Eagle Medium (DMEM) culture media (Media Core, Lerner Research Institute, Cleveland, OH) supplemented with 10% fetal bovine serum (FBS, GIBCO) and 1% penicillin-streptomycin in a 5% CO2-humidified chamber at 37°C. Cells were lentiviral transduced using the pLenti CMV V5-LUC Blast (w567-1) vector gifted by Eric Campeau (Addgene plasmid # 21474 ; http://n2t.net/addgene:21474 ; RRID:Addgene_21474) [37]. HEK293T cells obtained from the Lerner Research Institute, Cleveland, OH were grown in standard DMEM culture media conditions as outlined above. When 95% confluent, cells were transfected with the V5-Luciferase expression vector along with the pMD2.G and psPAX2 lentiviral packaging vectors. Transfection was performed using Lipofectamine 3000 Transfection Reagent (ThermoFisher Scientific, item no. L3000015) according to the manufacturer’s recommendations for lentiviral production and viral supernatant was harvested. Polyethylene glycol (PEG) solution was made by mixing 85 grams of PEG 8000 (MP Biomedicals, item no. 0219483901), 60 mL of 5M NaCl, 8 mL of 10x phosphate buffered saline (PBS) (Media Core, Lerner Research Institute, Cleveland, OH), and 32 mL of milliQ water, then brought up to a final volume of 200 mL by adding additional milliQ water. The solution was autoclaved, then placed on a rotator until cooled to room temperature. The PEG solution was added to virus containing supernatant in a 1:5 ratio by volume and allowed to sit for 24-48 hours at 4 degrees Celsius. This was then centrifuged at 1500 × g for 30 minutes, the viral pellet resuspended in PBS, and filtered through a 0.45 μm pore filter. A functional titer was performed using a 14-day treatment of 6 μg/μL blasticidin (ThermoFisher Scientific, item no. R21001). The selected pool of cells was used for all subsequent experiments including the generation of an ABHD6 knockout cell line as described below.

CRISPR-Cas9 genome editing was accomplished using methods previously described [38]. ABHD6 signal guide-RNAs (sgRNAs) were designed by an online tool (https://www.benchling.com/) and cloned into the Lenti-CRISPER v2 vector harboring a D10A nickase version of Cas9 (Cas9n) from Addgene [38]. Signal guides used for gene editing are as follows:

ABHD6-E5-Nick-5F 5’ CACCGAGGCTGGCTGACTCCTGCAG 3’,

ABHD6-E5-Nick-5R 3’ CTCCGACCGACTGAGGACGTCCAAA 5’,

ABHD6-E5-Nick-3F 5’ CACCGAGCAGGATGGATCTTGATG 3’,

ABHD6-E5-Nick-3R 3’ CTCGTCCTACCTAGAACTACCAAA 5’.

ABHD6Δ Huh7 cells were generated from the pLenti CMV V5-LUC Blast-transduced Huh7 cells following lentiviral transduction of the Lenti-CRISPER v2-Cas9 D10A-ABHD6 sgRNAs in similar fashion as described above for luciferase transduction. A functional titer was performed using a 7-day treatment of 3 μg/μL puromycin (Sigma, P8833-25MG). The selected pool of ABHD6Δ cells were validated by analyzing the expression of ABHD6 by western blot. Of note, the pool demonstrated virtually no ABHD6 expression therefore no single-cell / clonal isolation was performed. WT Huh7 cells were cultured alongside to account for passage number differences.

### Palmitic Acid-Induced Lipotoxicity

An 80 mM stock of palmitic acid (PA) (Santa Cruz Biotechnology, item no. sc-215881) in isopropanol was prepared, aliquoted, and stored at -80 degrees Celsius. On the day of treatment, PA stock was diluted 1:100 in 1% bovine serum albumin (Sigma, item no. A8806), serum-free DMEM media and sterilized through a 0.22 μm pore filter. For protein and RNA harvest, 2 million WT and Huh7 ABHD6Δ Huh7 cells were seeded into Falcon 60 mm tissue culture treated dishes (Corning, item no. 353002). After 24 hours, when cells were 70-80% confluent, media was replaced with 3 mL / plate of the 800 μM PA-containing media. After 24 hours of PA treatment, media was again replaced with 3 mL of fresh PA-containing media. Cells were harvested at baseline (just prior to the first PA treatment) and at various durations of palmitic acid treatment. Cells were harvested by scraping and further processing for protein and RNA extraction as described below.

### Cell Proliferation Assay

For cell proliferation assays, the Incucyte (Sartorius) live cell imager was used. 40,000 cells were seeded into a Costar 48-well tissue culture treated dish (Corning, item no. 3548), and gently mixed to ensure homogenous confluency, and the plate was positioned in the Incucyte chamber. After 24 hours from initial seeding, media was replaced with 200 μL / well of media containing treatment or vehicle, and image capture for cell proliferation analysis was started with the initial image used to determine “initial confluency.” Sixteen images were taken per well every 6 hours. The Basic Analyzer, “Standard” scan type, and “10x” magnification settings were used for image capture and analysis, with confluency normalized to initial confluency. Media containing treatment or vehicle was replaced every 24 hours. Treatments included 25 μM 18:1 BMP (S,S) (Avanti, item no. 857135P-5mg) suspended in 0.1% DMSO (Sigma, D2650), 2.5 ng/mL rapamycin (Santa Cruz Biotechnology, item no. sc-3504) suspended in 0.1% DMSO, or 30 μM chloroquine (Thermo Fisher Scientific, item no. P36239) suspended in 0.1% DMSO.

### Apoptosis Assay

For apoptosis assays, the Incucyte (Sartorius) live cell imager was used. 30,000 cells were seeded into a Falcon 96-well tissue culture treated white-wall clear-bottom plate (Corning, item no. 353377), and gently mixed to ensure homogenous confluency, and the plate was positioned in the Incucyte chamber. After 24 hours from initial seeding, cells were treated with 100 μL / well of 800 μM palmitic acid (PA) versus vehicle_PA_ (1% isopropanol (IPA) + 1% bovine serum albumin (BSA, Sigma, item no. A8806-5G)) with or without additional treatments (BMP, chloroquine, and rapamycin with corresponding vehicle as described above). After 24 hours of treatment, media was replaced and supplemented with the caspase-3/7 recognition motif DNA intercalating dye (Sartorius, item no. 4704) using a 1:1000 dye:media dilution and image capture was performed one-hour later. Five images were taken per well. The Basic Analyzer, “Adherent Cell-by-Cell” scan type, and “10x” magnification settings were used for image capture and analysis, with Red Object Count Per Image / Phase Area Confluence (1/Image / %) as the final readout value. A conversion factor was used to combine results from separate biologic replicates.

### Autophagy Assay with RFP-GFP-LC3 Reporter

Use of the Premo Autophagy Assays using the tandem RFP-GFP-LC3 reporter (ThermoFisher Scientific, item no. P36239) was done similar to previously described [39]. 20,000 cells in 300 μL of complete media were seeded into Falcon 8-well chambered cell culture slides (Corning, item no. 354118) and gently mixed to ensure homogenous confluency. After 24 hours from initial seeding, cells were transduced by adding 20 μL / well of the BacMam 2.0 baculovirus RFP-GFP-LC3 reagent. Twenty-four hours later, cells were treated with 150 μL / well of 800 μM palmitic acid (PA) versus vehicle (1% isopropanol (IPA) + 1% bovine serum albumin (BSA)). Twenty-four hours later, cells were fixed and taken for confocal microscopy imaging. Cell fixation was performed as follows: PBS and 5% FBS were warmed to 37 degrees Celsius. A stock 16% formaldehyde solution was diluted to 4% using the PBS + FBS solution. The wells were washed with the PBS + FBS solution once, then 500 μL / well of the created 4% formaldehyde solution was added and allowed to sit at room temperature for 20 minutes. The wells were then washed with PBS twice. The overlying chamber was removed. One drop of VECTASHIELD with DAPI reagent (Vector Laboratories, item no. H-1200) was placed over each well and the coverslip was mounted and secured. Slides were then imaged and analyzed as described below.

### LysoTracker Staining

60,000 cells were seeded into Nunc Lab-Tek Chambered Coverglass slides (ThermoFisher Scientific, item no. 155383) and gently mixed to ensure homogenous confluency. PA and corresponding vehicle were prepared as described above. After 24 hours from initial seeding, cells were treated with 400 μL / well of 800 μM palmitic acid (PA) versus vehicle (1% isopropanol (IPA) + 1% bovine serum albumin (BSA)). After 24 hours of treatment, media was replaced with 400 μL complete cell culture DMEM media containing 60 nM LysoTracker Deep Red (ThermoFisher Scientific, item no. L12492). Twenty-five minutes after replacement with LysoTracker-containing media, one drop of NucBlue DAPI stain (Invitrogen, item no. R37606) was added. Twenty minutes after the addition of NucBlue, the LysoTracker-containing media was replaced with 400 μL FluoroBrite DMEM media (ThermoFisher Scientific, item no. A1896701) and then taken for confocal microscopy imaging. Imaging and analysis were performed as described below.

### Confocal Microscopy and Image Analysis

Images from samples of both the autophagy and LysoTracker assays were acquired using a Leica TCS SP8-AOBS inverted confocal microscope equipped with 4 Hybrid Detectors, 405 nm Diode, Argon (488 nm), DPSS 561 nm, and HeNe 633 nm lasers (Leica Microsystems, GmbH, Wetzlar, Germany). In each experiment, images were previewed to identify samples with the strongest signal for each fluorophore for each assay respectively. Laser and detector settings were optimized using samples from each respective assay with the strongest signal for each independent fluorophore and saved. The saved settings were then applied to collect images from all samples for each respective assay. For each group a minimum of 5-6 fields at 40x were imaged containing ∼100 cells for analysis. Image analysis and quantitation was performed using Volocity 6.3 software (Quorum Technologies, Ontario, Canada). A separate library of the immunofluorescence images collected on the confocal was created in the Volocity software for each assay. The following steps outline the analysis of images using Volocity software, with a focus on ensuring consistent and accurate results. Images were processed for analysis to maintain uniformity by performing standardization of image settings for each batch of images. Image display settings were selected to minimize background signal utilizing the brightness and black level settings in the image tab. Of critical importance modifying the display settings does not change intensity values for the images, rather only the image display. The same image display settings for all the individual immunofluorescence channels were applied to each image for each respective assay to optimize visibility and consistency while avoiding over processing. A measurement protocol was then developed for each assay and their respective set of images to identify objects for each fluorophore. Objects were segmented using intensity and size filters. Once objects were segmented the compartmentalization tool was used to quantify the number of objects which overlap or are contained by another object for the different fluorophores. Cells were defined by presence of DAPI+ nuclei.

### Western Blots

Cell pellets and whole tissue homogenates were made from multiple tissues in a modified RIPA buffer, and Western blotting was conducted as previously described [40]. ImageJ software (National Institutes of Health, Bethesda, MD) was used for protein expression analysis. For the DMBA tumor development experiment, a previously generated rabbit polyclonal antibody against murine ABHD6 and anti-beta actin were used as previously described [12]. Information on the remainder of antibodies used may be found in Supplemental Table 1.

### Gene Expression Analyses by qPCR and Bulk RNA Sequencing

Tissue RNA extraction was performed as previously described [12]. Cell RNA extraction was performed using the RNeasy Plus Mini Kit (Qiagen) according to the manufacturer’s protocol. qPCR analyses were conducted as previously described [12]. The ΔΔ-CT method was used to determine relative mRNA expression. qPCR was conducted using the Applied Biosystems 7500 Real-Time PCR System. For RNA sequencing, cell pellet RNA extraction was performed using the RNeasy Plus Mini Kit (Qiagen) according to the manufacturer’s protocol. RNA samples were checked for quality and quantity using the Bio-analyzer (Agilent). RNA-SEQ libraries were generated using the Illumina mRNA TruSEQ Directional library kit and sequenced using an Illumina NovaSeq 6000 (both according to the Manufacturer’s instructions). RNA sequencing was performed by the University of Chicago Genomics Facility. Sequencing reads generated from the Illumina platform were assessed for quality and trimmed for adapter sequences using TrimGalore! v0.4.2 (Babraham Bioinformatics), a wrapper script for FastQC and cutadapt. Reads that passed quality control were then aligned to the human reference genome (GRCh38) using the STAR aligner v2.5.3 [41]. The alignment for the sequences were guided using the GENCODE annotation for GRCh38. The aligned reads were analyzed for differential expression using Cufflinks v2.2.1 [42], a RNASeq analysis package which reports the fragments per kilobase of exon per million fragments mapped (FPKM) for each gene. Differential analysis report was generated using the cuffdiff command performed in a pairwise manner for each group. Differential genes were identified using a significance cutoff of adjusted p-value (q-value) < 0.05. The genes were then subjected to gene set enrichment analysis (GenePattern, Broad Institute) to determine any relevant processes that may be differentially over represented for the conditions tested. Additional pathway analysis was performed using iPathwayGuide (Advaita Bio). Principle component analysis (PCA) plots were generated with ClustVis [43] using significantly differentially expressed genes. Prism v8.4.3 (GraphPad) was used to generate volcano plots, with significantly differentially expressed genes defined using an absolute value log2 fold change greater than 1 and adjusted p-value < 0.05. Prism v8.4.3 was used to generate heatmaps showing the top 25 up- and 25 down-regulated genes by log2 fold change. The RNAseq data included here have been deposited into the publicly-available Gene Expression Omniubus (GEO) portal with a temporary accession number of GSE255384.

### Targeted BMP Lipidomics

Cell pellets containing approximately 2 million cells transferred to 2 ml safe-lock tubes containing two 5 mm steel beads. Lipids were extracted utilizing a modified version described by Matyash et al. using 1 ml Methyl tert-butyl ether-methanol (3:1; v/v) containing 500 pmol butylated hydroxytoluene, 1% acetic acid, and 150 pmol internal standard (14:0-14:0 BMP, 17:0-17:0 PG; Avanti Polar Lipids, Alabaster, AL) per sample [44]. Subsequently, samples were homogenized by shaking on a Retsch Tissue-Lyser (Qiagen, Venlo, NL) for 2×10 sec (30hz). Extraction was performed under constant shaking on a thermomixer (1400 rpm) for 60 min at room temperature (RT). After the addition of 200 µl dH2O and further incubation for 20 min at RT, samples were centrifuged at 20,000 g for 10 min at RT on a tabletop Eppendorf centrifuge to establish phase separation. The upper organic phase was collected, dried under a stream of nitrogen and resolved in 200 µl methanol/2-propanol/H2O (6:3:1; v/v/v) for UPLC/MS analysis. Chromatographic separation was performed using an Agilent 1290 Infinity II UHPLC, equipped with a Kinetex C18 column (2.1×50 mm, 1.7µm; Phenomenex), a flow rate of 0.2 mL/min, an injection volume of 1 μL, column temperature of 50 °C, and a mobile phase gradient: (B %) from 37,5% to 100% B within 3 min, where solvent A was (methanol/H2O, 8/2, v/v;) and B was (2-propanol/H2O, 8/2, v/v;) both containing 10 mM ammonium acetate, 0,1% formic acid. An Agilent 6470 triple-quadrupole mass spectrometer with an Agilent Jet Stream ESI (Agilent Technologies, CA, USA) was used for detection. Fast polarity switching allowed for simultaneous analysis in ESI positive and negative in a single run. BMP and PG species were analyzed in dynamic multiple reaction monitoring mode, using the following transitions, in negative mode; [M-H]-to fatty acids anions for both classes, in positive mode; BMP: [MNH4]+ to [MG-H2O]+ of the respective MG, and, PG [MNH4]+ to [DAG-H2O]+ of the respective DAG. Unit resolution was used in both MS1 and MS2, the fragmentor voltage was set to 135 V for BMP, 300V for PG, and the collision energy was set to 25 V for BMP, 15V for PG. The ESI parameters were as follow: gas temp = 300°C; gas flow = 5 l/min; nebulizer = 30 psi; sheath gas temp = 400°C; capillary = 3500 V; in the positive mode: nozzle voltage = 0 V, while in the negative mode: nozzle voltage = 1000 V. The system was controlled by Agilent MassHunter Acquisition software version 10.1. Data processing was performed with Agilent MassHunter quantitative analysis software version 10.1 and Agilent MassHunter qualitative analysis software version 10.0. Data were normalized for recovery, extraction- and ionization efficacy by calculating analyte/internal standard ratios (arbitrary unit, AU) and expressed as AU/mg or g tissue weight/protein content.

### Targeted Monoacylglycerol (MAG) Lipidomics

Analysis of 1-MAG and 2-MAG species was done as described previously [16]. Briefly, total lipid was extracted from 5 million WT and ABHD6Δ Huh7 cells using Folch reagent. After lipid extraction, the lipids were dried under nitrogen stream and dissolved in 40µM chloroform. They were then loaded on silica gel thin layer chromatography (TLC) plates pre-coated with 2.3% boric acid. The TLC was developed in a solvent system (chloroform: acetone: acetic acid in a 60:40:1, v/v ratio) to separate 1-MAG and 2-MAG. The plates were exposed to iodine vapor and the spots corresponding to 1-MAG and 2-MAG were scraped, extracted and saponified. The released free fatty acids (FFAs) were extracted using Dole-Meinertz extraction procedure. The fatty acids in the n-heptane layer were dried under nitrogen stream and derivatized with phenacylbromide in order to be measured by reverse phase High Performance Liquid Chromatography (HPLC). A Zorbax Eclipse plus XDB analytical C18 column (4.6 × 250 mm; 5 μm; Agilent Technology) was used for the FFA species separation. FFAs were eluted using methanol/water/ACN/isopropanol (87.88/4.62/3.75/3.75) at a flow rate of 1.5 ml/min and detected at 242 and 254 nm. The quantification was done using FFA standard curves and by internal standard method. Of note, octanoic, decanoic, lauric, palmitoleic, arachidonic, and linoleic acid were included in this panel but their levels were undetectable.

### Hepatic Triglyceride Measurement

Extraction and measurement of hepatic triglycerides was performed as previously described [12].

### Statistics

Statistics were performed using Prism v8.4.3 (GraphPad). For the human HCC IHC analysis, results were described using medians with interquartile ranges for continuous variables, and total occurrences with percentages for categorical variables. For the remainder of the studies, univariate comparisons were made using unpaired two-tailed Fisher’s exact, Wilcoxon rank sum, or Student’s t-test. Body weight curve, tumor growth, and cell viability analyses were made using two-way analysis of variance (ANOVA).

## Results

### ABHD6 is Overexpressed in Human HCC Tumors

We first sought to determine expression levels of ABHD6 in human HCC samples. To do so, paraffin embedded sections from 83 patients who previously underwent partial liver resection for HCC of various etiology were analyzed using ABHD6-targeted immunohistochemistry (IHC) (**Fig.1A, 1B**). ABHD6 displayed a distinctive granular cytoplasmic perinuclear staining pattern (**Fig. 1A**), characteristic of a lysosomal staining pattern, consistent with previous data demonstrating ABHD6’s localization to the late endosome and lysosomal compartments in addition to nuclear and plasma membranes [22]. Lysosome-associated membrane protein (LAMP1)-directed IHC was performed and demonstrated a congruent staining pattern to ABHD6 (data not shown), further corroborating this localization scheme. Patient demographics and staining patterns from collected patient specimens are detailed in Supplemental Table 2. Patients were grouped by whether tumors demonstrated high or low ABHD6 expression levels as defined by granular cytoplasmic staining at 4x or 10x/20x magnification, respectively. No significant differences in patient demographics were observed. As expected, tumors with high ABHD6 expression also demonstrated more diffuse staining (81.67% vs 54.55%, p=0.021) defined as staining intensity observed in greater than 50% of visible tumor area. There was a trend toward increased lymphovascular invasion and recurrence in the higher expression group though analysis was limited by small sample size, implicating ABHD6 may play a role in cell invasion and migration. Strikingly, nearly all patients in both groups demonstrated increased ABHD6 IHC-staining in tumor cores compared to adjacent non-tumor liver tissue (**Fig. 1B, 1C**). To further investigate ABHD6’s protein expression profile in MASLD-related HCC, western blot on tumor and adjacent non-tumor tissue from 16 additional patients with HCC from exclusively MASLD etiology was performed. ABHD6 protein expression was higher in tumor cores compared to adjacent non-tumor liver tissue (**Fig. 1D, 1E**). Collectively, these data show ABHD6 is overexpressed in human HCC tumor cores from various etiologies, including MASLD.

**Figure 1.**
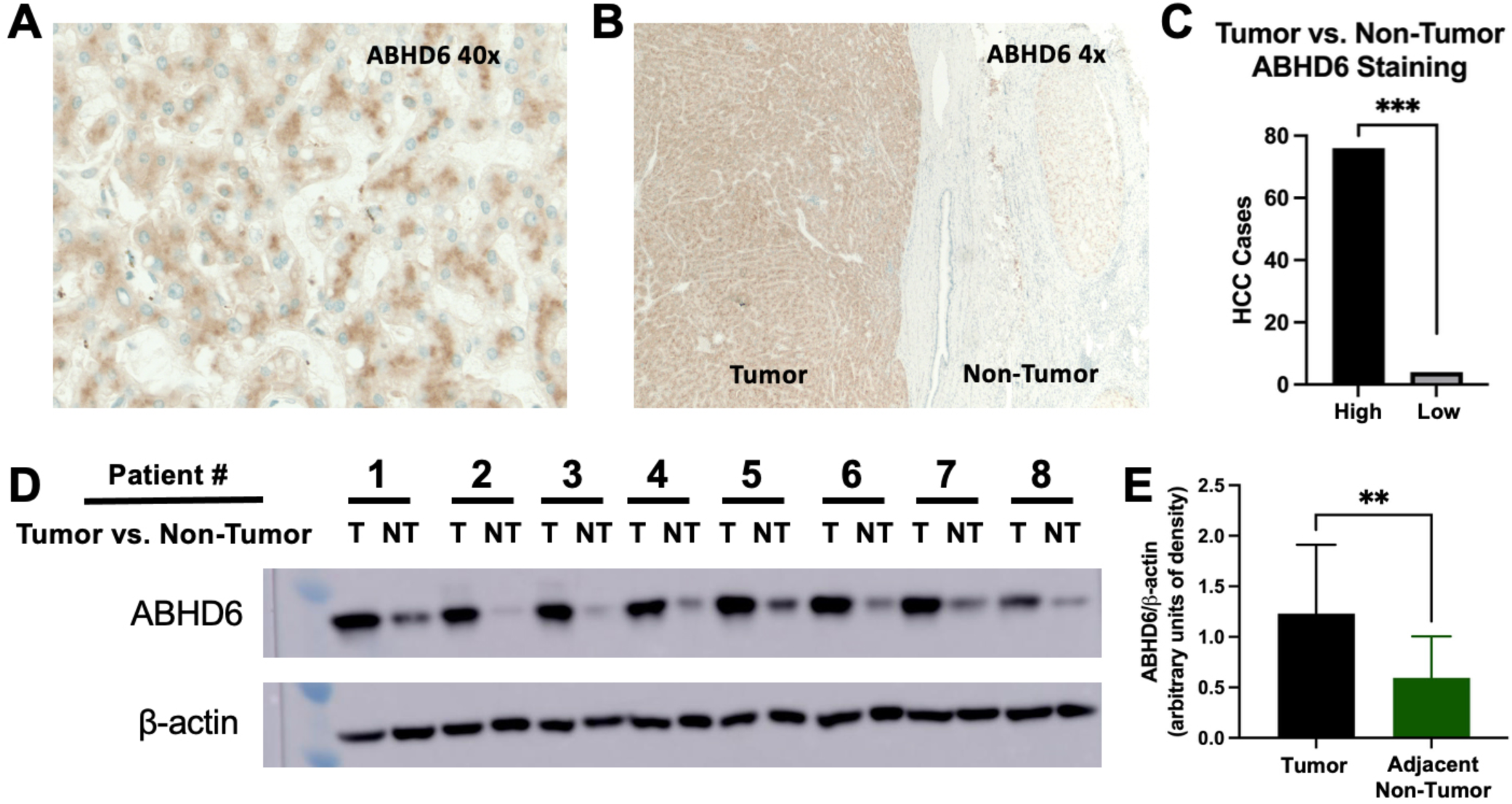
ABHD6 is Upregulated in Human HCC. **(A)** Immunohistochemical (IHC) targeting ABHD6 was performed on patient tumor specimens from a variety of etiologies, though predominantly viral. At 40x magnification, ABHD6-targeted IHC demonstrates finely granular cytoplasmic perinuclear staining in HCC tumor cells. **(B)** Representative image of ABHD6-targeted IHC at 4x magnification demonstrates strong cytoplasmic granular staining in the HCC tumor core (left). The adjacent non-tumorous cirrhotic liver parenchyma (right) reveals faint and patchy cytoplasmic granular staining. **(C)** IHC slides were collectively analyzed and patient tumors were found to have ABHD6 overexpression in tumor cores compared to adjacent non-tumor. Chi-square test. n=80. **(D)** Western blot analysis was performed on patient tumor specimens from exclusively MASLD etiology. Representative western blot shows ABHD6 protein expression is upregulated MASLD-related HCC tumor vs adjacent non-tumor liver tissue. **(E)** Western blots were collectively analyzed, and the resultant densitometry analysis shown. Paired t-test. n=16 patient tumor – non-tumor pairs (32 samples total). Graphs displayed as mean +/- SD. *p<0.05, **p<0.01, ***p<0.001. T: tumor; NT: non-tumor.

### ABHD6 Inhibition Reduces HCC Development in an Obesity/MASLD-Driven Mouse Model

Next, we sought to investigate the role of ABHD6 in an HCC preclinical animal model. We used the obesity-driven DMBA mouse model [34], where pups are given a one-time treatment of the carcinogen DMBA. Tumor development requires a second liver insult in the form of liver steatosis achieved through long-term high fat diet feeding, as chow fed mice do not develop liver tumors. This model’s dependence on high fat diet feeding make it particularly relevant to the study of MASLD-related HCC [34,45]. After weaning, mice were randomized to receive either a peripherally-acting (without penetration into the central nervous system) ABHD6-targeting antisense oligonucleotide (ASO) or a scrambled non-targeting control ASO to investigate the effects of ABHD6 inhibition on tumor development. ABHD6-targeting ASOs effectively decreased ABHD6 mRNA and protein expression (**Fig. 2A**) and resulted in drastically reduced tumor burden, as measured by the number of grossly visible tumors at time of necropsy and by blinded pathologist scoring (**Fig. 2B-D**). A trend towards decreased lung tumor formation in the ABHD6-targeting ASO group was also observed (80% vs 20%, p=0.206) (Supplemental Fig. 1). Additionally, ABHD6 inhibition led to a decrease in body weight, liver-to-body weight ratio, and liver triglycerides (**Fig. 2E-G**), ultimately blunting the development of high fat diet induced metabolic syndrome. qPCR demonstrated a reduction in genes involved in the lipogenesis and cholesterol synthesis including acetyl CoA-carboxylase 1 (*Acaca*), fatty acid synthase (*Fasn*), sterol regulatory element-binding protein-1c (*Srebf1c*), *Srebf2*, and 3-hydroxy-3-methylglutaryl-coenzyme A reductase (*Hmgcr*), as well as Adhesion G Protein-Coupled Receptor E1 (*Adgre1*) encoding the murine macrophage marker F4/80 (**Fig. 2H**). Overall, these results demonstrate ABHD6 inhibition alters liver lipid metabolism and reduces tumor development in a mouse model of obesity-driven HCC.

**Figure 2.**
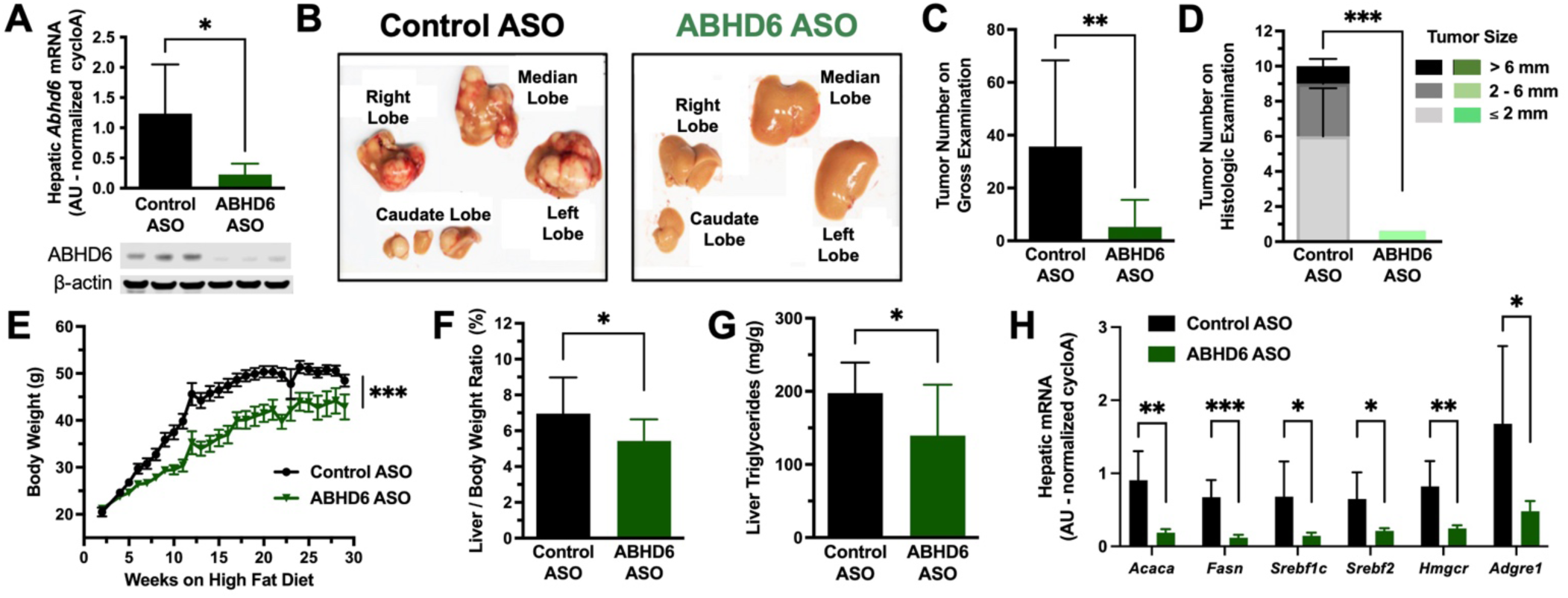
ABHD6 Knockdown Prevents Liver Tumor Formation in a Mouse Model of Obesity-Driven HCC. C57BL/6 pups were treated with the carcinogen dimethylbenz[a]anthracene (DMBA) at 3 days of age. At 21-days of age mice were weaned to a high fat diet, and randomized to receive an ABHD6-targeting antisense oligonucleotide (ASO) or a control scrambled ASO. Mice were maintained on HFD for 30 weeks until necropsy. **(A)** ABHD6-targeting ASOs effectively reduced ABHD6 liver mRNA, unpaired t-test, and protein expression, representative image. **(B)** Representative image of ASO-mediated inhibition of ABHD6 on tumor burden. **(C)** ABHD6 inhibition significantly reduced tumor burden assessed by grossly visualized total tumor number per liver by two independent data image reviewers, Wilcoxon rank sum test, as well as by **(D)** histological analysis with tumor size sub-classification, two-way ANOVA. **(E)** ABHD6 inhibition significantly reduced weight gain, two-way ANOVA, as well as **(F)** liver-to-body weight at time of necropsy, unpaired t-test, and **(G)** liver triglycerides, unpaired t-test. **(H)** ABHD6 inhibition significantly altered hepatic mRNA expression of multiple metabolically relevant genes as measured by qPCR, multiple unpaired t-tests. n=10-15 with exceptions: n=3 for western blot, n=5 for histological analysis, n=6 for qPCR. Graphs displayed as mean +/- SD with the exception of panel (E) displayed as mean +/- SEM. *p<0.05, **p<0.01, ***p<0.001. AU: arbitrary units; Acaca: acetyl CoA-carboxylase 1; Fasn: fatty acid synthase; Srebf1c: sterol regulatory element-binding protein-1c; Hmgcr: 3-hydroxy-3-methylglutaryl-coenzyme A reductase; Adgre1: Adhesion G Protein-Coupled Receptor E1.

### ABHD6 Inhibition Reduces HCC Progression in an Obesity/MASLD-Driven Mouse Model

While the previous experiment demonstrated ABHD6 inhibition prevents tumor *development* in the obesity-driven DMBA model, we wanted to explore if AHBD6 inhibition would have an effect if initiated after tumor formation i.e. if it affects tumor *progression*. This distinction is especially important in determining if AHBD6 harbors a direct antitumorigenic effect, as ABHD6 blunted the development of metabolic syndrome (**Fig. 2E-G**) which is a critical component for tumor formation in this model. Therefore, we initiated ASO treatment after tumors were formed and visualized by ultrasound (**Fig. 3A**), rather than at 3 weeks of age as in the prior experiment. At 25 weeks of age, we performed liver ultrasound weekly. After tumors were well visualized, mice were matched based on tumor size and randomized to receive either an ABHD6-targeting ASO or a scrambled non-targeting control ASO to investigate the effects on ABHD6 inhibition on the progression of liver tumors. Tumors were well delineated on ultrasound imaging (**Fig. 3A**) and weekly measurements were taken to track tumor growth until endpoint. The ABHD6-targeting ASO was effective in reducing both tumor and non-tumor ABHD6 protein expression (**Fig. 3B**). ABHD6 inhibition significantly reduced tumor progression over a median of 7 weeks of treatment (**Fig. 3C**), supporting its role as a therapeutic target in HCC. In addition, we observed a significant decrease in body weight with ABHD6 inhibition over the treatment course (**Fig. 3D**). The observation that ABHD6 inhibition blunts tumor progression in this clinically translational experimental design, with drug treatment initiated after tumor formation, provides strong support that ABHD6 inhibition displays a direct antitumorigenic effect.

**Figure 3.**
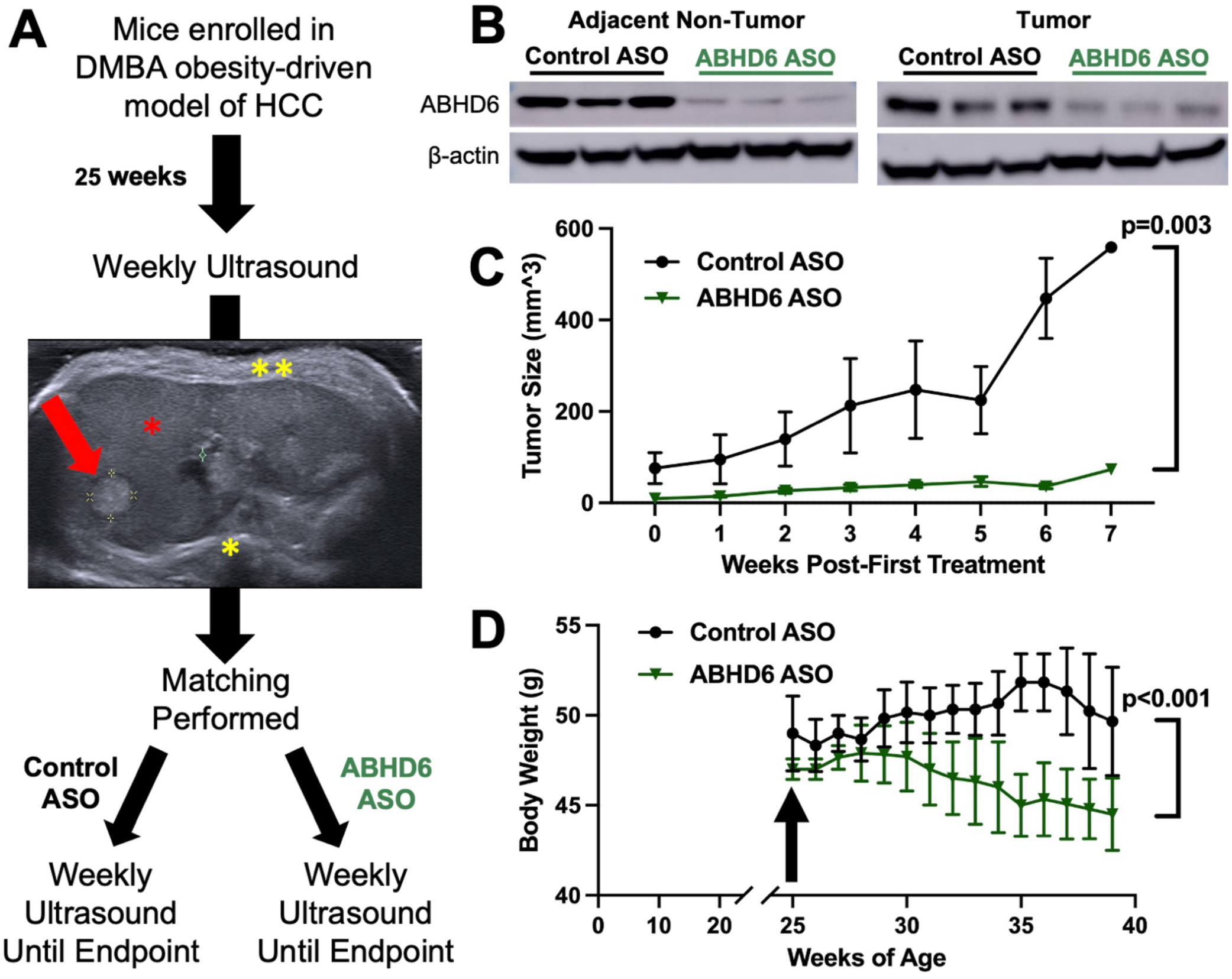
ABHD6 Knockdown Initiated After Tumor Formation Reduces Tumor Progression in a Mouse Model of Obesity-Driven HCC. C57BL/6 pups were treated with the carcinogen dimethylbenz[a]anthracene (DMBA) at 3 days of age. At 21-days of age mice were weaned to a high fat diet. **(A)** Beginning at 25-weeks of age, mice were imaged with ultrasound weekly to assess tumor burden. Mice were matched based on tumor burden and randomized to receive an ABHD6-targeting antisense oligonucleotide (ASO) or a control scrambled ASO, and weekly ultrasound imaging was performed. A representative ultrasound image of a tumor is shown, red arrow: tumor, red asterisk: non-tumor liver parenchyma, single yellow asterisk: spine, double yellow asterisk: abdominal wall. Ultrasound well delineated tumor borders, and was used to serially assess tumor size. **(B)** ABHD6-targeting ASOs effectively reduced ABHD6 protein expression in tumors as well as non-tumorous adjacent liver tissue (representative western blot shown). **(C)** ABHD6 inhibition reduced tumor progression measured by serial tumor size from time of first treatment. **(D)** Mice treated with ABHD6-targeting ASOs (black arrow = time of first ASO treatment) demonstrated a significant reduction in body weight compared to control. Two-way ANOVA, n=4-6. Graphs displayed as mean +/- SEM.

### Knockout of ABHD6 Increases BMP Lipids but Not MAG Lipids in a Human Hepatoma Cell Line

To study the effect of ABHD6 on lipid and tumor biology we created an ABHD6 genetic knockout line in Huh7 human hepatoma cells. Using a double-nickase CRISPR-Cas9, ABHD6 knockout (ABHD6Δ) Huh7 cells were generated and knockout confirmed via western blot (**Fig. 4A**, Supplemental Fig. 2). Next, we measured MAG and BMP lipids, two known lipid substrates of ABHD6 that had been previously shown to be relevant to cellular pathways of glucose and lipid metabolism. There were no differences in total MAG lipid levels, or in diverse molecular species of MAGs (**Fig.4B**, Supplemental Fig. 3). In contrast, ABHD6Δ cells had significantly increased total BMP lipids (**Fig. 4C**), with notable increases observed in several subspecies (**Fig. 4D**). Interestingly, BMP lipids increased cell proliferation in both ABHD6Δ and wild type (WT) cells, though no genotype-specific differences were observed (Supplemental Fig. 4). Taken together, these data show the successful achievement of functional ABHD6 knockout in the Huh7 human hepatoma cell line which served as a useful tool in the following experimental models.

**Figure 4.**
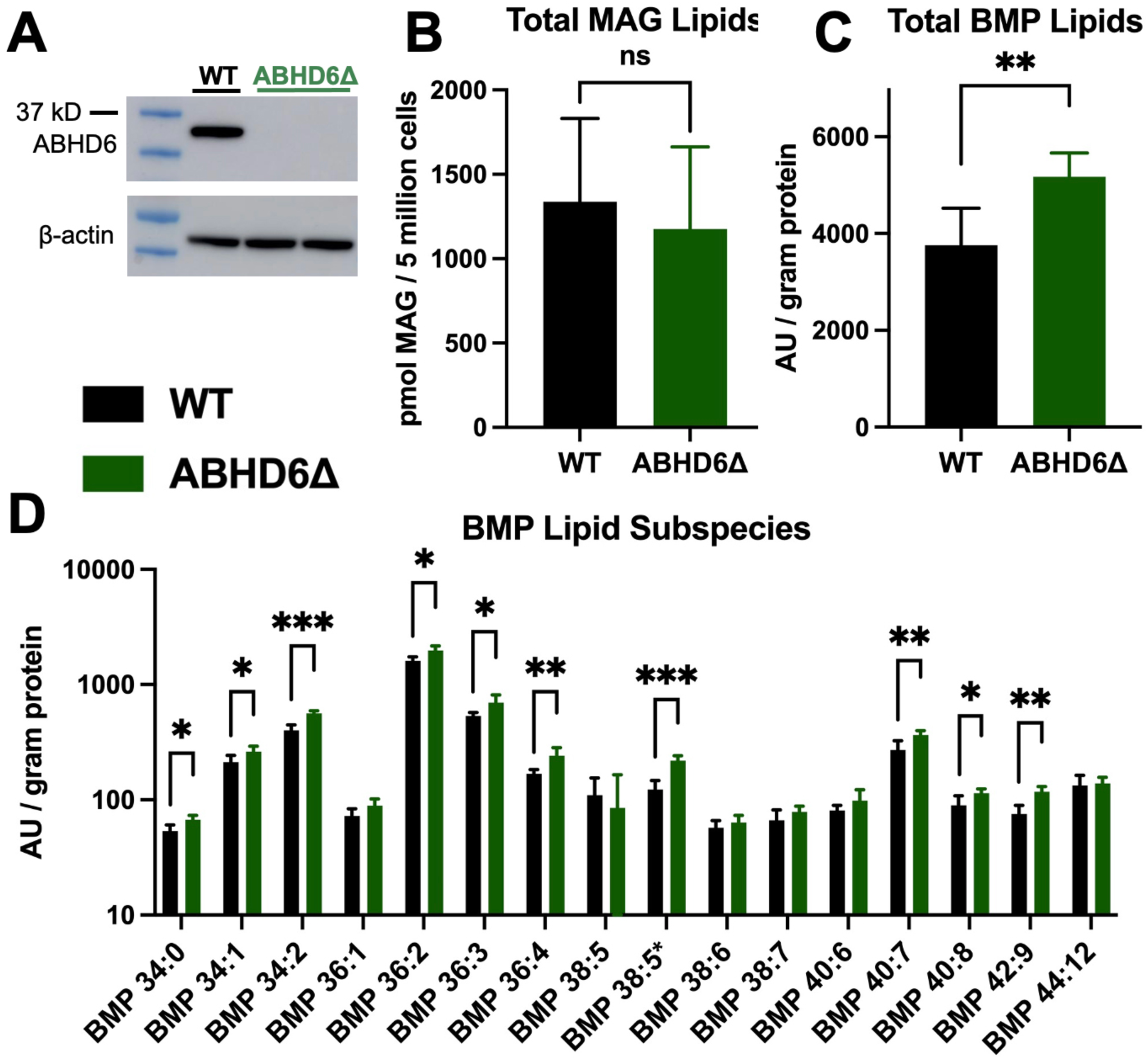
ABHD6 Knockout Alters the Lipidomic Profile in a Human Hepatoma Cell Line. Huh7 wild type (WT) and ABHD6Δ cells were cultured under basal media conditions for 24 hours, then harvested for lipidomic analysis. **(A)** Western blot analysis confirmed the absence of ABHD6 protein expression in ABHD6Δ cells (representative western blot shown). **(B)** No difference was seen in total monoacylglycerol (MAG) lipids between groups; however, both **(C)** total bis(monoacylglycerol)phosphates (BMP) lipids and **(D)** BMP lipid subspecies were found to be significantly different. Multiple unpaired t-tests, n=4-5. Graphs displayed as mean +/- SD. *p<0.05, **p<0.01, ***p<0.001, ns: not significant. AU: arbitrary units; BMP 38:5 (18:1-20:4); BMP 38:5* (18:2-20:3).

### Genetic Knockout of ABHD6 Reduces Tumor Engraftment in an Orthotopic Xenograft Model

To investigate the role of ABHD6 on tumor development in the human hepatoma Huh7 cell line, we utilized an orthotopic xenograft model with luciferase-transduced WT and ABHD6Δ cells. Briefly, as a survival surgery, a laparotomy was performed and either WT or ABHD6Δ cells were injected in two separate locations within the liver. Following surgery, mice were fed a high fat diet to recapitulate the MASLD component of hepatic lipid overload, and bioluminescence imaging was used to serially track engraftment until necropsy, which was five weeks after surgery (**Fig. 5A**). Ultrasound imaging was performed just prior to necropsy to both help determine tumor engraftment and quantify tumor size (ultrasound image not shown). Tumors were well-encapsulated spherical masses that resembled human HCC both grossly and histologically (**Fig. 5B, 5C**), and genetic knockout was maintained in ABHD6Δ tumors over the duration of the experiment (**Fig. 5D**). To determine rates of tumor engraftment, it was assumed there were two tumors possible per mouse given that two tumor cell liver parenchymal injections were performed per mouse, hence the total number of tumors possible per group = 2 x # of mice per group. ABHD6 knockout resulted in a trend toward reduced tumor engraftment by bioluminescence signal (**Fig. 5E**), and produced significantly less tumors at time of necropsy (61.1% vs 25%, p=0.034) (**Fig. 5F**). We observed no difference in tumor size, though this was limited by the small sample size of the knockout group (**Fig. 5G**). In line with the obesity-driven DMBA mouse model results, these results demonstrate ABHD6 knockout reduces tumor engraftment in a xenograft model using a human hepatoma cell line.

**Figure 5.**
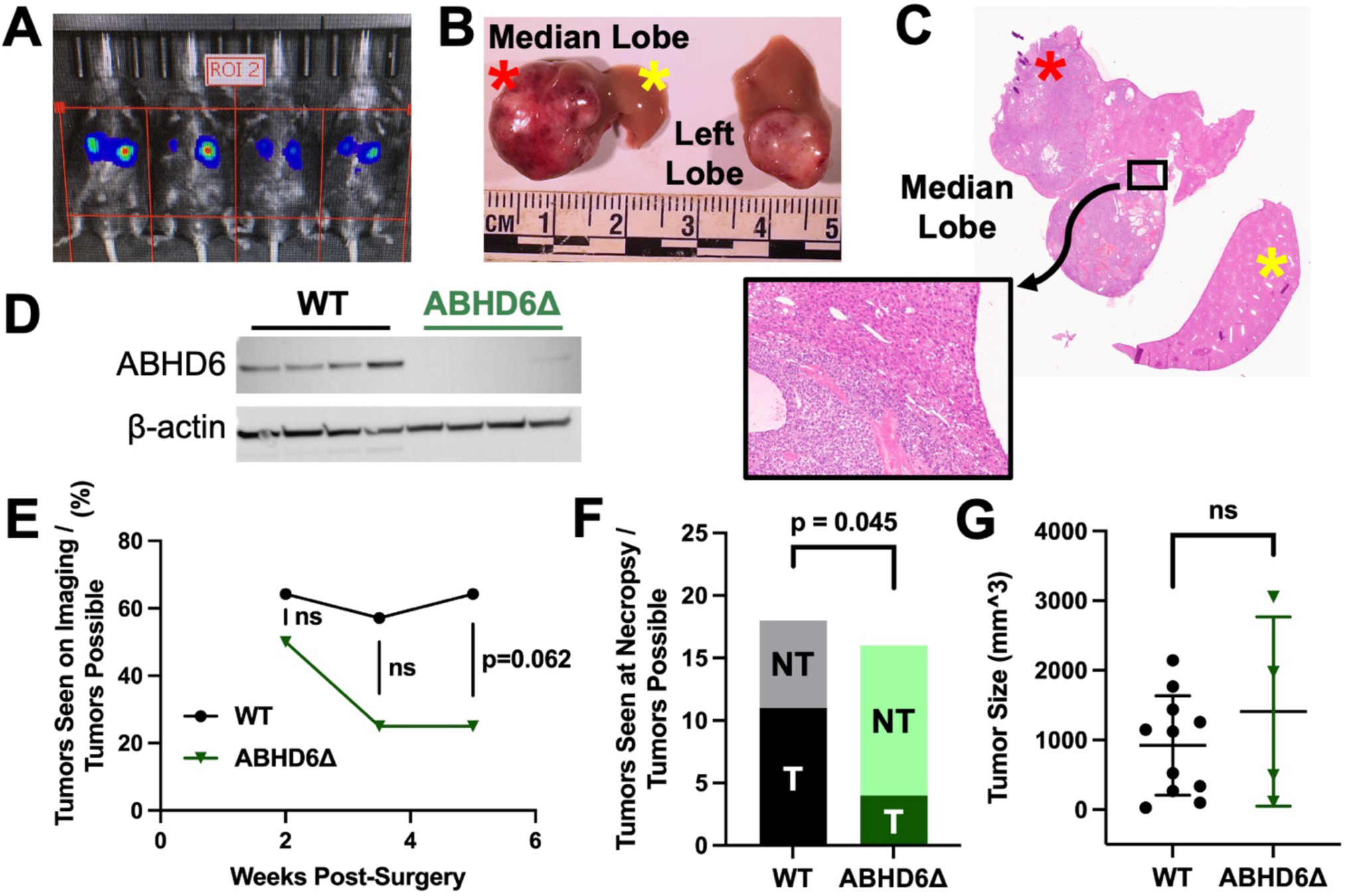
ABHD6 Knockout Reduces Tumor Engraftment in an Orthotopic Xenograft Model of HCC. 15-week-old C57BL6 Rag2-knockout mice were placed on a high fat diet for 2 weeks prior to surgery. An orthotopic xenograft was performed via laparotomy and subcapsular injection of 1 million luciferase-transduced Huh7 wild type (WT) or ABHD6Δ cells into the right side of the median liver lobe and another 1 million cells into the inferior portion of the left liver lobe. Mice were euthanized at 5 weeks post-surgery. **(A)** Luciferase-based bioluminescence imaging (signal represents Huh7 tumor cell luciferase activity) and ultrasound imaging (not shown) were performed to serially determine tumor engraftment and size. **(B)** Pictured is the median and left liver lobes at time of necropsy, red asterisk: right side of median liver lobe, yellow asterisk: left side of median liver lobe. On gross inspection tumors appeared solitary, encapsulated, and hyper-vascularized. **(C)** Pictured is H&E staining of the tumorous median liver lobe only i.e. no left liver lobe shown from panel (B), with corresponding red and yellow asterisks. Tumor H&E staining demonstrates similar morphology and growth to human HCC. Higher magnification demonstrates the non-tumor (upper right)-tumor (bottom left) junction. **(D)** ABHD6 knockout was maintained throughout the duration of the experiment as evidenced by tumor western blot analysis at time of necropsy (representative western blot shown). **(E)** The ABHD6Δ group trended toward displaying reduced rates of bioluminescence signal over the experiment (final timepoint p=0.062 using multiple Mann-Whitney U tests, p=0.070 using Kaplen Meier curve analysis [not shown in graph]). **(F)** Tumor engraftment was determined by completion ultrasound prior to necropsy as well as by gross examination of the liver at time of necropsy. A significant reduction of tumor engraftment was observed in the ABHD6Δ group compared to WT, 11/18 (61.1%) vs 4/16 (25%), p= 0.045, Fisher’s exact test. **(G)** No significant difference was observed in tumor size by ultrasound imaging, unpaired t-test. n=8 and n=9 mice in the WT and ABHD6Δ groups, respectively (n=16 and n=18 potential sites for engraftment in the WT and ABHD6Δ groups, respectively), with the following exceptions: n=4 tumors for western blot analysis, n=6-7 mice for bioluminescence analysis. Graphs displayed as mean +/- SD. ns: not significant.

### ABHD6 Knockout Alters the Transcriptomic Profile of a Human Hepatoma Cell Line in a Cell Culture Model of Lipotoxicity

We used an established cell culture model of palmitic acid (PA)-induced lipotoxicity and autophagy to study the effects of ABHD6 deletion in Huh7 cells [25]. WT and ABHD6Δ Huh7 cells were subjected to 24 hours of PA treatment then used for RNA sequencing analysis. PCA analysis demonstrated distinct clustering between genotypes (**Fig. 6A**). Volcano plot analysis (**Fig. 6B**) and heatmap plots of hierarchically clustered differentially expressed genes (**Fig. 6C**) exhibited clear differences between WT and ABHD6Δ cells. After the 24 hours of palmitic acid treatment, pathway differences between genotypes were focused on cell stress responses, notably the phosphoinositide 3-kinase-protein kinase B (PI3K-Akt) signaling pathway and autophagy (**Fig. 6D**). Since apoptosis may contribute to transcriptional differences in cellular stress responses, we performed an apoptosis assay in WT and ABHD6Δ Huh7 treated with PA. Additionally, we performed the apoptosis assay with BMP lipid treatment given our earlier data demonstrating increased BMP lipids in ABHD6Δ cells (**Fig. 4**) as well as genotype-independent increased cell proliferation with BMP treatment (Supplemental Fig. 4). There were no genotype-dependent differences in apoptosis with or without the addition of BMP lipids, though BMP treatment did reduce levels of apoptosis in a genotype-independent manner (Supplemental Fig. 5). Under basal cell culture conditions i.e. void of PA-induced lipotoxicity, gene expression differences were also observed though all significantly altered gene pathways were lipid metabolism-related rather than cell stress response pathways and included the hypoxia inducible factor 1-alpha (HIF1-alpha), peroxisome proliferator-activated receptor (PPAR), and AMP-activated protein kinase (AMPK) signaling pathways (Supplemental Fig. 6). Taken together, these findings suggest ABHD6 deletion in Huh7 cells produces transcriptomic changes, specifically lipotoxicity-induced alterations in autophagy and other cell stress related pathways.

**Figure 6.**
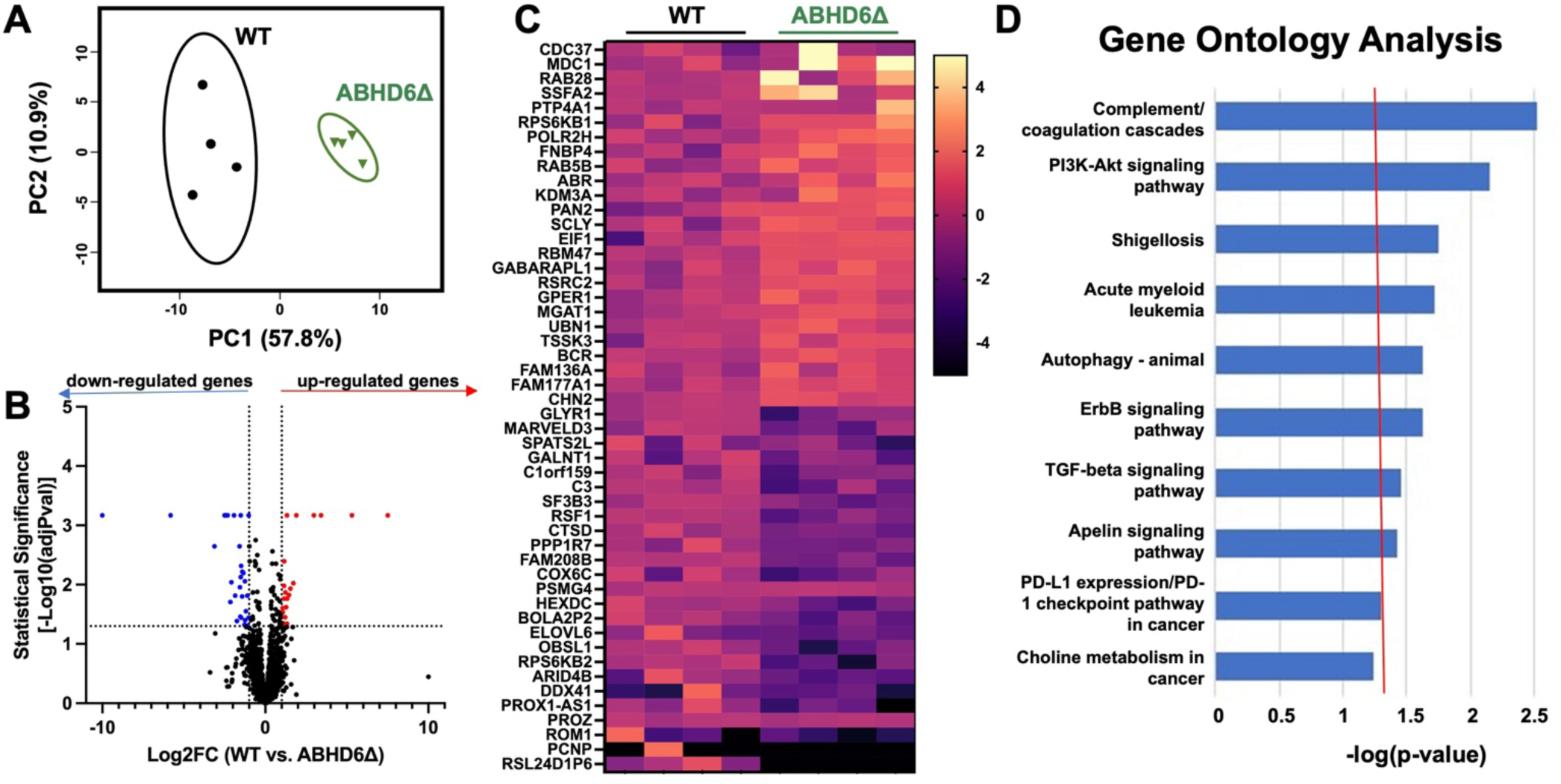
ABHD6 Knockout Alters Lipotoxicity-Induced Autophagy and Cell-Stress Relevant Pathways in a Human Hepatoma Cell Line. Huh7 wild type (WT) and ABHD6Δ cells were treated with 800 μM palmitic acid (PA) for 24 hours and harvested for RNAseq analysis. **(A)** Principal component analysis showed distinct clustering of WT and ABHD6Δ cells. **(B)** Volcano plot of gene expression changes (n = 4; genes with an absolute value log2 fold change greater than 1 and adjusted p-value < 0.05 were considered significantly differentially expressed), **(C)** and heat map of row-normalized expression for the top 50 differentially expressed genes demonstrate strong gene expression changes between groups. **(D)** Gene ontology analysis was notable for differences multiple cell-stress relevant pathways including autophagy and the PI3K-AKT signaling pathway.

### Knockout of ABHD6 Alters Autophagy and Lysosomal Activity of a Human Hepatoma Cell Line in a Cell Culture Model of Lipotoxicity

Since our pathway analysis revealed altered autophagy signaling in our previous experiment, we further investigated autosomal and lysosomal function in WT and ABHD6Δ Huh7 cells treated with PA for up to 48 hours (**Fig. 7A-D**, Supplemental Fig. 7). We observed significant differences in the autosomal proteins microtubule-associated protein 1A/1B-light chain 3 (LC3), ubiquitin-binding protein p62, and mammalian target of rapamycin (mTOR) as well as in lysosomal-associated membrane protein 2 (LAMP2). Next, we assessed autophagic flux using the RFP-GFP-LC3 gene reporter (**Fig. 7E-G**, Supplemental Fig. 8). WT and ABHD6Δ Huh7 cells were transduced with the gene reporter then treated with PA for 24 hours and imaged using confocal microscopy. ABHD6Δ cells displayed higher RFP signal with a trend toward increased GFP fluorescence as well, suggesting increased engagement of autosomal pathways. Autophagic flux evidenced by RFP/GFP signal appeared similar between genotypes (data not shown). To relate these differences to the endosomal-autosomal-lysosomal system, the LysoTracker fluorescent dye was used (**Fig. 7H-I**, Supplemental Fig. 9). WT and ABHD6Δ Huh7 cells were treated with PA or vehicle alone for 24 hours then labeled with the basic LysoTracker dye which indiscriminately stains all acidic compartments of a cell, predominantly late-endosomes/endolysosomes, autolysosomes, and lysosomes. Interestingly, ABHD6Δ cells displayed lower signal than WT cells when subjected to either PA treatment or vehicle alone, which is in contrast to the RFP-GFP-LC3 gene reporter experiment which only looked at the autophagosome and autolysosome compartments. Together, this may suggest a shuttling of lysosomes toward autophagy as a result of ABHD6 knockout. Of note, the differences seen in fluorescent signal measured by “area of fluorescent puncta” in both the RFP-GFP-LC3 gene reporter and LysoTracker experiments were secondary to differences in the number of puncta/objects seen and not due to differences in the size of those puncta/objects (data not shown), suggesting against morphologic differences from ABHD6 knockout as being the cause. Lastly, we used rapamycin and chloroquine which serve to activate or inhibit autophagy, respectively, to study proliferation and lipotoxicity-induced apoptosis (Supplemental Fig. 10). With the exception of chloroquine treatment in the apoptosis assay, no genotype-specific differences were observed. As a whole, these data suggest that ABHD6 knockout alters autosomal and lysosomal pathways in Huh7 cells.

**Figure 7.**
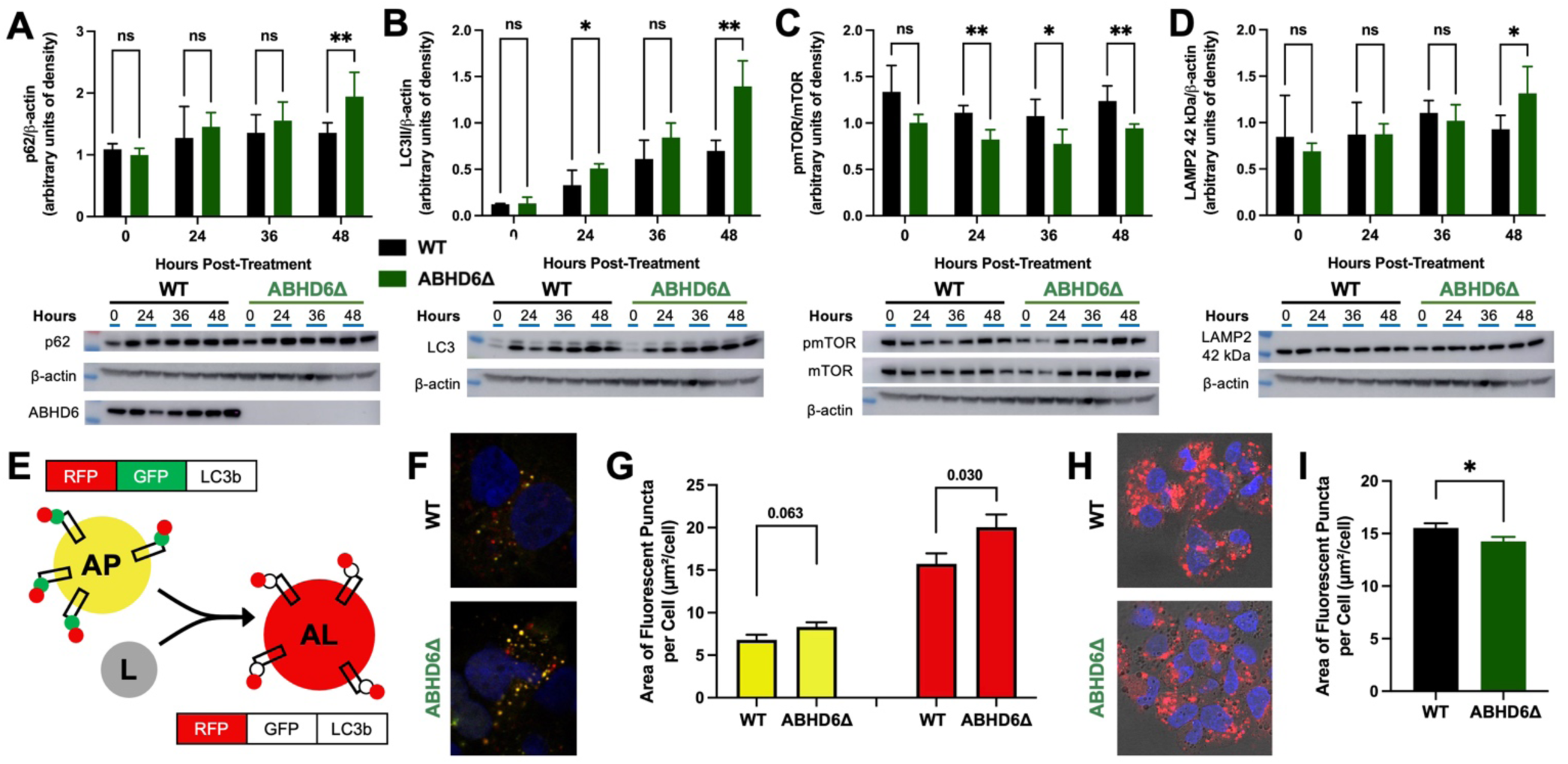
ABHD6 Knockout Alters Autophagy and Lysosomal Activity in a Human Hepatoma Cell Line Subjected to Palmitic Acid-Mediated Lipotoxicity. **(A-D)** Huh7 wild type (WT) and ABHD6Δ cells were treated with 800 μM palmitic acid (PA) for up to 48 hours. Cells were harvested for western blot analysis. Densitometry analysis (above) and representative western blots (below) demonstrate significant changes in p62, LC3II, pmTOR, and LAMP2 at various timepoints. ABHD6 knockout was confirmed (representative western blot shown in panel A). Multiple Mann-Whitney tests, n=2. Experiment performed in biological triplicate (total n=6), results combined and resultant graphs displayed as mean +/- SD. (**E)** Huh7 WT and ABHD6Δ cells were treated with 800 μM PA for 24 hours, and the RFP-GFP-LC3 reporter was used to study autophagic flux. Displayed is a schematic of RFP-GFP-LC3 reporter. LC3 localizes to the autophagosome (AP), and dual RFP and GFP fluorescence results in a yellow signal. Upon fusion of the autophagosome and lysosome (L) to yield the acidic autolysosome (AL), a red signal results due to the acid-sensitive nature of GFP. **(F)** Representative images (DAPI used for nuclear staining) and **(G)** quantification of GFP (left) and RFP (right) performed demonstrating significantly higher RFP signal (p=0.030) in ABHD6Δ cells signifying increased autophagosome plus autolysosome formation. **(H)** Huh7 WT and ABHD6Δ cells were treated with 800 μM PA for 24 hours. LysoTracker Deep Red staining was performed. Representative images and **(I)** quantification demonstrate decreased signal in ABHD6Δ cells. The RFP-GFP-LC3 reporter assay and LysoTracker staining experiments were each analyzed using an unpaired t-test, n=2-4, experiments performed in biological duplicate, results combined and resultant graphs displayed as mean +/- SEM. *p<0.05, **p<0.01, ***p<0.001.

### Small Molecule Inhibition of ABHD6 Reduces Tumor Progression in a Clinically-Relevant Orthotopic Xenograft Model

In an effort to investigate the role of ABHD6 inhibition in the aforementioned orthotopic xenograft model using a more clinically translational design, we employed small molecule-mediated inhibition on WT tumors after tumors were formed and visualized on ultrasound imaging (**Fig. 8A**). To make clear, in contrary to the previously performed xenograft experiment in Figure 5, all tumors in this experiment were generated using WT cells and the mode of ABHD6 modulation was via treatment with a small molecule inhibitor rather than the use of CRISPR-Cas9 mediated knockout cells. An orthotopic xenograft surgery using only WT Huh7 cells was performed, with mice maintained on high fat diet perioperatively to recapitulate the MASLD component. One week after surgery, mice were randomized to receive KT203, a peripheral dual mouse-/human-targeting irreversible small molecule inhibitor of ABHD6, or vehicle alone. We imaged liver tumors by ultrasound at 21 days, 26 days, and 31 days postoperatively to track tumor growth. Tumors were well delineated on ultrasound imaging and measurements were taken for volume calculation (**Fig. 8B**). The administration of the ABHD6-inhibitor, KT203, led to a significant reduction in tumor progression (**Fig. 8C**). Notably, tumor engraftment was identical between groups (81% vs 81%). This clinically relevant experimental design using a small molecule inhibitor on well-established liver tumors provides strong support for ABHD6 inhibition as a therapeutic option in HCC.

**Figure 8.**
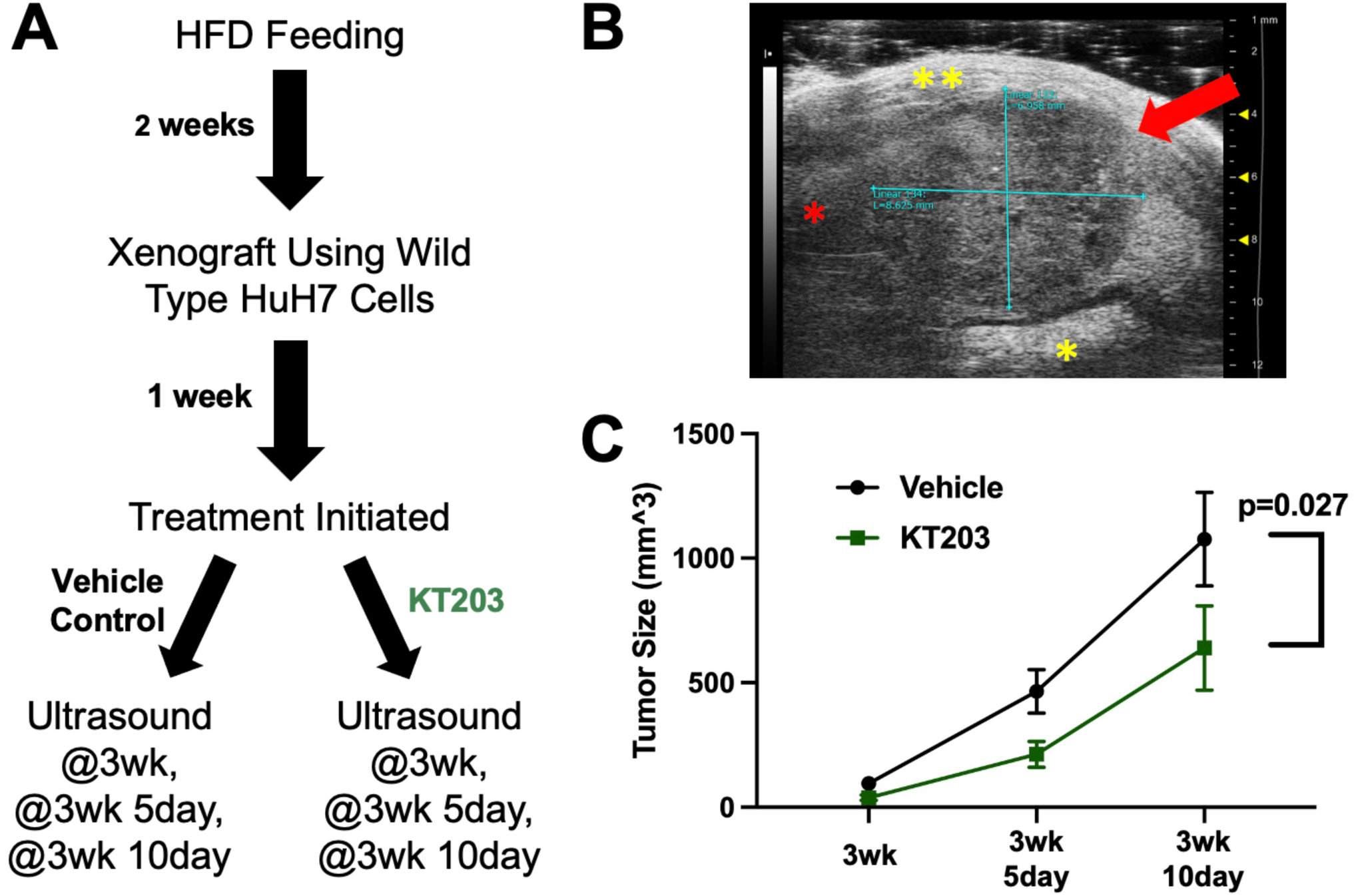
Small Molecule Inhibition of ABHD6 Reduces Tumor Progression in in an Orthotopic Xenograft Model of HCC. **(A)** 15-week-old C57BL/6 Rag2-knockout mice were placed on a high fat diet for 2 weeks prior to surgery. An orthotopic xenograft was performed via laparotomy and subcapsular injection of 1 million luciferase-transduced Huh7 wild type (WT) cells in the right side of the median liver lobe and in the inferior portion of the left liver lobe. One-week post-surgery mice were randomized to receive the dual mouse-/human-targeting ABHD6 small molecule inhibitor, KT203, versus vehicle (18:1:1 saline:PEG40:ethanol) alone daily via intraperitoneal injection. Mice were euthanized at 5 weeks post-surgery. **(B)** An example of ultrasound imaging of tumors is shown, red arrow: tumor, red asterisk: non-tumor liver parenchyma, single yellow asterisk: stomach, double yellow asterisk: abdominal wall. Ultrasound well delineated tumor borders, and was used to serially assess tumor size. **(C)** KT203 treatment significantly reduced tumor progression, p=0.027, two-way ANOVA, n=13. Graph displayed as mean +/- SEM.

## Discussion

MASLD rates have risen rapidly over the past few decades and it has become a leading cause of HCC [3–6]. Mechanistic overlaps are being explored in efforts to achieve both MASLD treatment and HCC chemoprevention using a single therapeutic [8–10]. ABHD6 was previously shown to be a critical regulator of metabolic syndrome [12], and was also found to be involved in the pathogenesis of various cancers [19,20]. Herein, we show that ABHD6 plays a role in the development and progression of HCC. Major findings of this work include: 1) ABHD6 was upregulated in human HCC tumor cores compared to adjacent non-tumorous liver, 2) ASO-directed knockdown of ABHD6 prevented HCC development and blunted tumor progression in an obesity-driven mouse model of HCC, 3) genetic knockout of ABHD6 in a human hepatoma cell line altered relevant cell stress pathways including autophagy and lysosomal activity, and reduced tumor engraftment in an orthotopic xenograft model, and 4) use of a small molecule inhibitor of ABHD6 reduced HCC progression in an orthotopic xenograft model. Collectively, these results provide support for ABHD6 as a potential therapeutic target in HCC.

Our group has previously demonstrated the pivotal role of ABHD6 in metabolic syndrome and steatotic liver disease [12], which we confirmed in this study (**Fig. 2F-I**), but little was known about ABHD6’s function in cancer. A recent large screen of over 700 drugs of interest found two ABHD6 inhibitors to be the leading candidates for preventing pancreatic cancer metastasis, and results were confirmed using a subcutaneous xenograft mouse model [19]. More recently, Tang et al. found ABHD6 inhibition prevented non-small cell lung cancer (NSCLC) growth and metastasis in a xenograft model through monoacylglycerol (MAG)-mediated suppression of epithelial-to-mesenchymal transition (EMT) [20]. Given its established role in liver lipid metabolism, we sought to determine if ABHD6 played a role in MASLD-related HCC, which has not been previously investigated.

Importantly, we performed studies in an obesity-driven mouse model of HCC to recapitulate MASLD-related HCC [34]. We first used antisense oligonucleotides (ASOs), which were employed to achieve a peripheral inhibition of ABHD6 in our prior studies [12], and found liver tumor development was markedly reduced (**Fig. 2**). Interestingly, there was also a trend in reduced lung tumor formation (Supplemental Fig. 1), providing support for the aforementioned findings on ABHD6 in NSCLC. Our subsequent experiments generated more clinically relevant data by demonstrating that ABHD6 ASO treatment in mice with pre-formed liver tumors reduced tumor progression (**Fig. 3**), further supporting ABHD6 could be a therapeutic target in HCC. Since high fat diet-induced hepatotoxicity is required for liver tumor development in the DMBA model, we postulated that ABHD6 knockdown may work indirectly through liver lipid metabolism modulation rather than via an independent anti-tumor mechanism to explain our findings. To address this, we generated a CRISPR-Cas9 genetic knockout of ABHD6 in the human hepatoma cell line Huh7. Using an *in vitro* lipotoxicity model with palmitic acid (PA) [25], ABHD6 altered cell stress-driven pathways such as PI3K signaling and autophagy which play vital roles in tumorigenesis and progression (**Fig. 6**). We confirmed our findings in an orthotopic xenograft model where ABHD6Δ Huh7 cells displayed significantly lower rates of engraftment, providing strong evidence for an antitumorigenic role in HCC. Lastly, using a peripheral dual mouse-/human-targeting small molecule inhibitor of ABHD6 that was administered beginning 1 week after xenograft (**Fig. 8**), we found that ABHD6 inhibition reduced tumor progression. This finding in a clinically relevant and translational experimental model provides strong data for the therapeutic efficacy of ABHD6 inhibition in HCC.

Substrates of ABHD6 include 2-arachidonylglycerol (2-AG) [11], MAGs [13,21], lysophospholipids [12], and bis(monoacylglycerol)phosphates (BMPs) [22]. While early research focused on ABHD6’s contribution to 2-arachidonylglycerol metabolism and endocannabinoid signaling in the central nervous system, we previously demonstrated that ABHD6 plays a relatively minor role on these signaling pathways in the liver [12]. Similarly, ABHD6 only had a slight impact on hepatic MAG hydrolysis [22], likely due to high hepatic MAGL expression. In contrast, hepatic lysophospholipids, especially lysophosphatidylglycerol (LPG) species, are heavily regulated by ABHD6 [12], and their phosphatidylglycerol (PG) substrate lipids were found to be significantly increased in ABHD6Δ Huh7 cells (data not shown). However, given the intriguing relationship between late endosomal/lysosomal pathways to MASLD and cancer [25–33], we focused our efforts on BMP lipids in this paper. ABHD6 localizes to late-endosomes/lysosomes and is responsible for the high majority of hepatic BMP lipid turnover [22]. ABHD6 knockout mice display higher plasma BMP lipids [23]. BMP lipids are present in intralumenal vesicles of late-endosomes/lysosomes where they play a role in lysosomal function and lipid sorting [24]. Hepatic and plasma BMP lipid levels are increased in lysosomal storage diseases and after long-term high fat diet feeding, and positively correlate with MASLD severity in human patients [23]. In line with these previous findings, we show ABHD6 follows a lysosomal staining pattern in human HCC specimens (**Fig.1**) and genetic knockout produces increased cellular BMP lipids in ABHD6Δ Huh7 cells (**Fig. 4**).

Lysosomes are intimately involved in lipid metabolism [28,46]. Autophagy, the process of cellular “self-eating,” relies on lysosomes for successful fusion and formation of the autolysosome [26]. Autophagy also plays a leading role in cellular lipid metabolism with mTOR and AMPK, both master nutrient sensors of the cell, regulating its activity. Additionally, autophagy functions in the cellular stress response and cancer-relevant pathways [29–33]. This made the use of a cell culture model of PA-induced lipotoxicity attractive [25], with existing literature as well as data within this manuscript demonstrating its ability to induce autophagy and lysosomal processes [39]. ABHD6Δ Huh7 cells demonstrated increased microtubule-associated protein 1A/1B-light chain 3 (LC3) II expression but there was no difference in LC3II/1 ratio (**Fig. 7B**, Supplemental Fig. 7), suggesting a possible increase in autophagy activation but without increased autophagic flux i.e. without increased fusion of the autophagosome with the autolysosome. Increased ubiquitin-binding protein p62 expression seen in ABHD6Δ cells (**Fig. 7A**) may support this notion, though alternatively could represent a dysfunction in autophagy. Using confocal microscopy, ABHD6Δ cells demonstrated increased autophagosome plus autolysosome compartments (**Fig. 7G**) though decreased late-endosome/endolysosome plus autolysosome plus lysosome compartments (**Fig. 7I**). Taken together, the confocal microscopy data may suggest a shuttling of lysosomes toward autophagy in ABHD6Δ cells. It remains unclear whether these changes occur because of enhanced or impaired autophagy in ABHD6Δ cells and further investigation is ongoing. Notably, no changes in cell proliferation or in lipotoxicity-induced apoptosis were observed between genotypes with or without the supplementation of rapamycin or chloroquine with the single exception of chloroquine treatment in the lipotoxicity-induced apoptosis experiment (Supplemental Fig. 4, Supplemental Fig. 10). Additionally, despite elevated intracellular BMP lipids in ABHD6Δ Huh7 cells, the ability of BNP lipids to confer increased cell proliferation (Supplemental Fig. 4) and reduced lipotoxicity-induced apoptosis (Supplemental Fig. 5) occurred independent of genotype. The lack of genotype-specific differences observed *in vitro* is in stark contrast to the robust *in vivo* data encompassing two mouse models and multiple methods to inhibit or knockdown ABHD6. The contrast seen between *in vivo* and *in vitro* studies as well as the inability for BMP lipids and established modulators of autophagy (rapamycin, chloroquine) to produce genotype-specific differences *in vitro* suggest a multifaceted effect of ABHD6 knockout on carcinogenesis and tumorigenicity. It is worth noting that ABHD6 may play a role in the early stages of tumor formation given the drop-off of ABHD6Δ xenograft tumors seen by bioluminescence imaging (**Fig. 5E**) and the decreased tumor engraftment seen at endpoint whereas KT203 treatment which was initiated one-week after surgery had no effect on engraftment rate. Also, prior studies found ABHD6 inhibition reduces the EMT phenotype which has high crossover with tumor engraftment [19,20], though one study found this to be through modulation of MAG lipids which are not strongly impacted by hepatic ABHD6 [20].

In conclusion, our work highlights the therapeutic potential of targeting ABHD6 in HCC. Our rigorous approach included multiple mouse models of HCC and employed several methods to modulate ABHD6 including ASOs, CRISPR-Cas9 gene editing, and small molecule inhibition. While both of our HCC protocols utilized a high fat diet, the DMBA and xenograft models are distinct with one involving carcinogenic administration and the latter a deficient immune system.

Modes of inhibition vary, with likely differences in the biodistribution of ASOs and KT203 and their degree of inhibition. Also, the ubiquitous expression of ABHD6 and lipid metabolism differences between mice and humans further complicate extrapolation. Although ABHD6 knockout alters autophagy and liposomal activity, the exact mechanisms and overall effect on cellular processes is unclear. While further investigation is needed, ABHD6’s role in metabolic syndrome and liver lipid metabolism make it an attractive target in MASLD. With steadily rising rates of MASLD-related HCC, a therapeutic target common to both would be highly advantageous serving in both treatment and chemoprevention. Collectively, we provide support that ABHD6 may be a novel therapy for both the treatment of MASLD and chemoprevention of HCC and warrants further investigation.

## Author Contributions

*Conceptualization*, D.O. and J.M.B.; *Methodology*, D.O., W.J.M., K.K.F., V.V., I.R., R.B., D.J.S., L.J.O., A.L.B., S.M., D.F., S.C., R.C.S., C.F., C.N., A.C.B., A.J.H., P.P., R.N.H., D.B., R.Z., Y.H.L., S.R.M.M., M.P., R.G.L., A.E.M., O.Z., S.D., Z.L., D.S.A., F.A., J.D.L, and J.M.B.; *Investigation*, D.O., W.J.M., K.K.F., V.V., I.R., R.B., D.J.S., L.J.O., A.L.B., S.M., D.F., S.C., R.C.S., C.F., C.N., A.C.B., A.J.H., P.P., R.N.H., D.B., R.Z., Y.H.L., S.R.M.M., M.P., R.G.L., A.E.M., O.Z., Z.L., D.S.A., F.A., J.D.L, and J.M.B.; *Validation*, D.O., W.J.M., K.K.F., V.V., I.R., R.B., and J.M.B.; *Formal analysis*, D.O., W.J.M., K.K.F., V.V., I.R., R.B., D.J.S., L.J.O., A.L.B., S.M., D.F., S.C., R.C.S., C.F., C.N., A.C.B., A.J.H., P.P., R.N.H., D.B., R.Z., Y.H.L., S.R.M.M., M.P., R.G.L., A.E.M., O.Z., Z.L., D.S.A., F.A., J.D.L, and J.M.B.; *Writing – original draft*, D.O. and J.M.B.; *Writing – reviewing and editing*, D.O., W.J.M., K.K.F., V.V., I.R., R.B., D.J.S., L.J.O., A.L.B., S.M., D.F., S.C., R.C.S., C.F., C.N., A.C.B., A.J.H., P.P., R.N.H., D.B., R.Z., Y.H.L., S.R.M.M., M.P., R.G.L., A.E.M., O.Z., S.D., Z.L., D.S.A., F.A., J.D.L, and J.M.B.; *Funding acquisition*, J.M.B.; *Supervision*, J.M.B.

## Acknowledgements

This work was supported in part by National Institutes of Health grants R01 DK130227 (J.M.B.), P01 HL147823 (J.M.B.), P50 AA024333 (J.M.B.), T32 GM088088 (L.J.O), American Heart Association Postdoctoral Fellowships 15POST2535000 (R.C.S.) and 17POST3285000 (R.N.H.). This study was also supported by funds from the Canadian Institutes of Health Research (#143308 to MP and SRMM). We received internal funding support from the Cleveland Clinic Liver Tumor Center of Excellence (COE). WJM is supported by a Cleveland Clinic Global Center for Pathogen and Human Health Research Postdoctoral Fellowship provided by the Infection Biology Program of the Cleveland Clinic. Use of the Vevo 2100 (Visual Sonics) ultrasound was supported by NIH small instrumentation grant 1S10ODO21561 and use of the IVIS Spectrum In Vivo Imaging System was supported by NIH shared instrument grant S10OD018205. We are grateful to John Peterson and Ajay Zalavadia in the Cleveland Clinic Imaging Core for help with imaging quantification of autophagy flux. We are also thankful to Dr. Kimberly Such for consultation and collaboration of animal studies requiring coordinated veterinary care. We are grateful to Dr. R. Matthew Walsh and Dr. Jeremy Lipman for financial support provided to Dr. Danny Orabi during his joint time in surgical training at the Cleveland Clinic and as a student in the Molecular Medicine Ph.D. program of Case Western Reserve University.

## Note

Supplementary data for this article is available at … (http://)

