## Supplemental Information for "The Lipid Hydrolase ABHD6 is a Therapeutic Target in Metabolic Dysfunction-Associated Steatotic Liver Disease (MASLD)-Related Hepatocellular Carcinoma"

**Supplemental Table 1. Antibody Information.**

| Antibody | Vendor | Working Dilution | Product # |
|---|---|---|---|
| **ABHD6 (D3C8N)** | Cell Signaling | 1:1,000 | 97573 |
| **p62** | Fitzgerald | 1:10,000 | 20R-PP001 |
| **LC3A/B (D3U4C)** | Cell Signaling | 1:1,000 | 12741 |
| **mTOR (7C10)** | Cell Signaling | 1:1,000 | 2983 |
| **phospho-mTOR (Ser2448) (D9C2)** | Cell Signaling | 1:1,000 | 5536 |
| **p70 S6K1 (49D7)** | Cell Signaling | 1:1,000 | 2708 |
| **phospho-p70 S6K1 (Thr389) (108D2)** | Cell Signaling | 1:1,000 | 9234 |
| **AMPKα** | Cell Signaling | 1:1,000 | 2532 |
| **Phospho-AMPKα (Thr172) (40H9)** | Cell Signaling | 1:1,000 | 2535 |
| **LAMP1 (D2D11)** | Cell Signaling | 1:1,000 | 9091 |
| **LAMP2 (D5C2P)** | Cell Signaling | 1:1,000 | 49067 |
| **TFEB (D2O7D)** | Cell Signaling | 1:1,000 | 37785 |
| **ATP6V1A** | GeneTex | 1:1,000 | GTX110815 |
| **ATP6V1B2** | GeneTex | 1:1,000 | GTX110783 |
| **β-Actin, HRP-Linked** | Proteintech | 1:5,000 | HRP-60008 |
| **Rabbit IgG, HRP-linked** | Cell Signaling | 1:5,000 | 7074 |
| **Guinea Pig IgG, HRP-linked** | Abcam | 1:5,000 | ab6908 |

| | Strong ABHD6 Expression (n=61) | Low-to-Moderate ABHD6 Expression (n=22) | p-value |
|---|---|---|---|
| **Age (years)** | 57.0 [53.0:64.0] | 58.0 [53.0:63.0] | 0.868 |
| **Gender** | | | 0.136 |
| **male** | 51 (83.61%) | 15 (68.18%) | |
| **female** | 10 (16.39%) | 7 (31.82%) | |
| **Etiology** | | | 0.282 |
| **NAFLD** | 5 (8.2%) | 2 (9.09%) | |
| **Alcohol** | 9 (14.75%) | 0 (0%) | |
| **Viral Hepatitis B** | 3 (4.92%) | 1 (4.55%) | |
| **Viral Hepatitis C** | 34 (55.74%) | 13 (59.09%) | |
| **Cryptogenic** | 2 (3.28%) | 3 (13.64%) | |
| **PBC** | 3 (4.92%) | 0 (0%) | |
| **Other/Multiple** | 5 (8.2%) | 3 (13.64%) | |
| **Area of Tumor with Expression** | | | **0.021** |
| **negative** | 0 (0%) | 0 (0%) | |
| **5-50% (focal)** | 11 (18.33%) | 10 (45.45%) | |
| **>50% (diffuse)** | 49 (81.67%) | 12 (54.55%) | |
| **n/a** | 1 | 0 | |
| **Background Cirrhosis** | 59 (96.72%) | 21 (95.45%) | 1.000 |
| **Grade** | | | 0.306 |
| **well differentiated** | 18 (31.03%) | 8 (40%) | |
| **moderately differentiated** | 32 (55.17%) | 10 (50%) | |
| **poorly differentiated** | 8 (13.79%) | 2 (10%) | |
| **n/a** | 3 | 2 | |
| **Lymphovascular Invasion** | | | 0.214 |
| **yes** | 15 (24.59%) | 2 (9.52%) | |
| **no** | 46 (75.41%) | 19 (90.48%) | |
| **n/a** | 0 | 1 | |
| **Recurrence** | 6 (9.84%) | 0 (0%) | 0.334 |
| **Follow Up (months)** | 61.0 [31.0:114.0] | 96.5 [51.3:129.3] | 0.208 |

**Supplemental Table 2. Patient Characteristics of HCC Specimens used for ABHD6 Immunohistochemistry.** Immunohistochemical (IHC) targeting ABHD6 was performed on patient tumor specimens from a variety of etiologies. Tumors were grouped by degree of ABHD6 expression. Strong versus low/moderate ABHD6 expression was defined as demonstrating granular cytoplasmic staining at 4x or 10x/20x magnification, respectively. P-values were calculated by Wilcoxon rank sum test or one-way analysis of variance (ANOVA). PBC: primary biliary cholangitis.

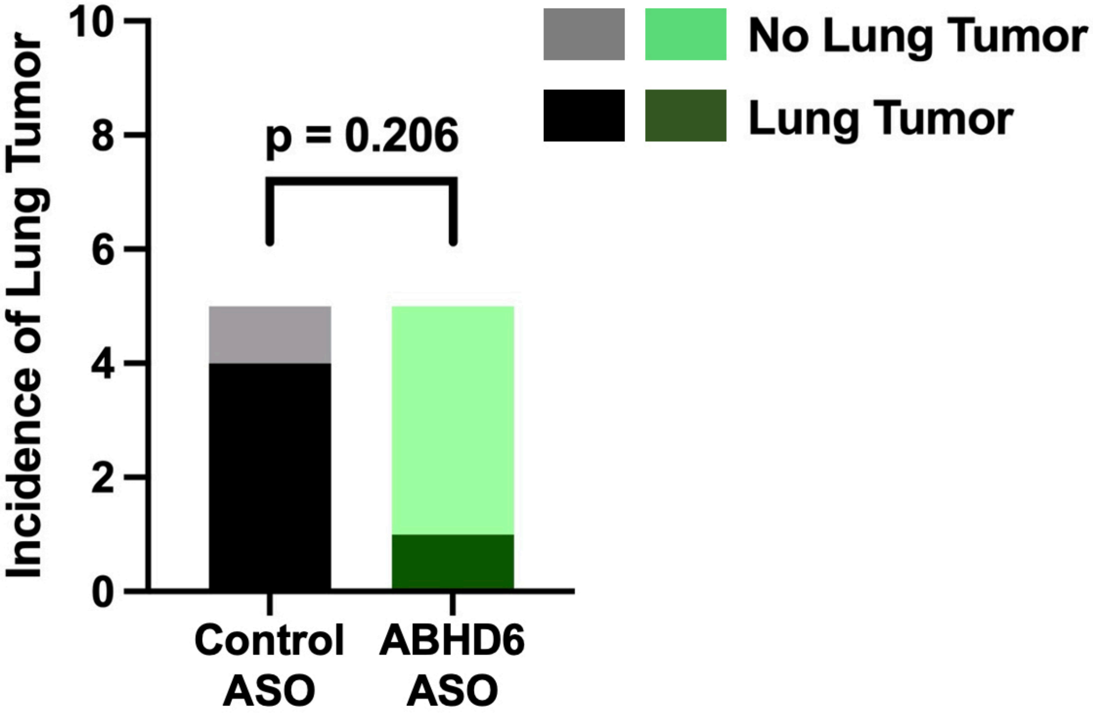


**Supplemental Figure 1. ABHD6 Knockdown Prevents Lung Tumor Formation in a Mouse Model of Obesity-Driven HCC.** C57BL/6 pups were treated with the carcinogen dimethylbenz[a]anthracene (DMBA) at 3 days of age. At 21-days of age mice were weaned to a high fat diet, and randomized to receive an ABHD6-targeting antisense oligonucleotide (ASO) or a control scrambled ASO. Mice were maintained on HFD for 30 weeks until necropsy. Lung tumor burden assessment by histological analysis demonstrated ABHD6 inhibition trended toward decreased lung tumor formation (1/5 vs 4/5, p=0.206), Fisher's exact test, n=5.

agttttaatatcgtcattctctttggcccctgcagGAG
TCAGCCAGCCTGAAAGAGCAGGATGGATC
TTGATGTGGTTAACATGTTTGTGATTGCGG
GCGGCACGCTGGCCATCCCAATCCTGGCA
TTTGTGGCTTCATTTCTTCTGTGGCCTTCA
GCACTGATAAGAATCTATTATTGgtaagccag
ttttatcattgatgttttcaagag

**Supplemental Figure 2. Generation of Double-Nickase CRISPR-Cas9 Mediated Huh7 ABHD6Δ Human Hepatoma Cells.** ABHD6 sgRNAs were designed by an online tool (https://www.benchling.com/) and cloned into the Lenti-CRISPER v2 vector (Addgene (Ran et al., 2013) with D10A nickase version of Cas9 (Cas9n)). The desired genomic deletion was centered around the predicting mRNA ATG start codon (underlined) at the beginning of exon 3. After transduction of the Lenti-CRISPER v2-Cas9 D10A-ABHD6 sgRNAs in Huh7 cells and 7 days of puromycin selection, the resultant pool of cells (ABHD6Δ cells) demonstrated virtually no ABHD6 protein expression (as shown in Fig. 4A). The hg38_knownGene_ENST00000478253.6 genomic sequence was used (UCSC Genome Browser). Intron: lowercase letters; exon: capital letters; orange + red letters: ‘NGG’ sequence of protospacer adjacent motif (PAM) site; blue letters: targeted signal guide RNAs.

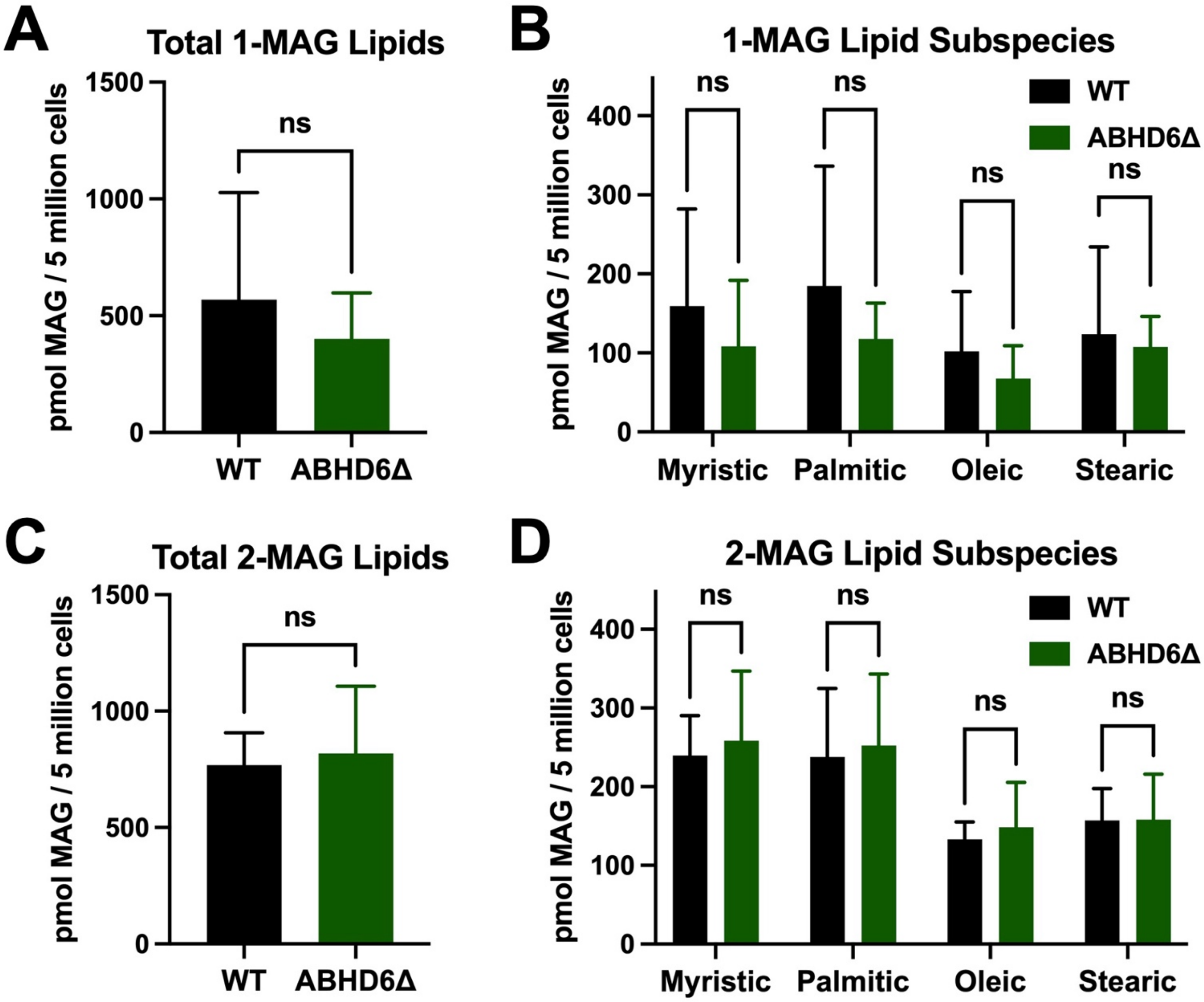


**Supplemental Figure 3. ABHD6 Knockout Does Not Alter MAG Lipids in a Human Hepatoma Cell Line.** Huh7 wild type (WT) and ABHD6Δ cells were cultured under basal media conditions for 24 hours, then harvested for lipidomic analysis. No significant differences were seen in **(A)** total and **(B)** respective 1-monoacylglycerol (MAG) subspecies, nor **(C)** total and **(D)** respective 2-MAG subspecies. Unpaired t-test, n=4. Graphs displayed as mean +/- SD. ns: not significant.

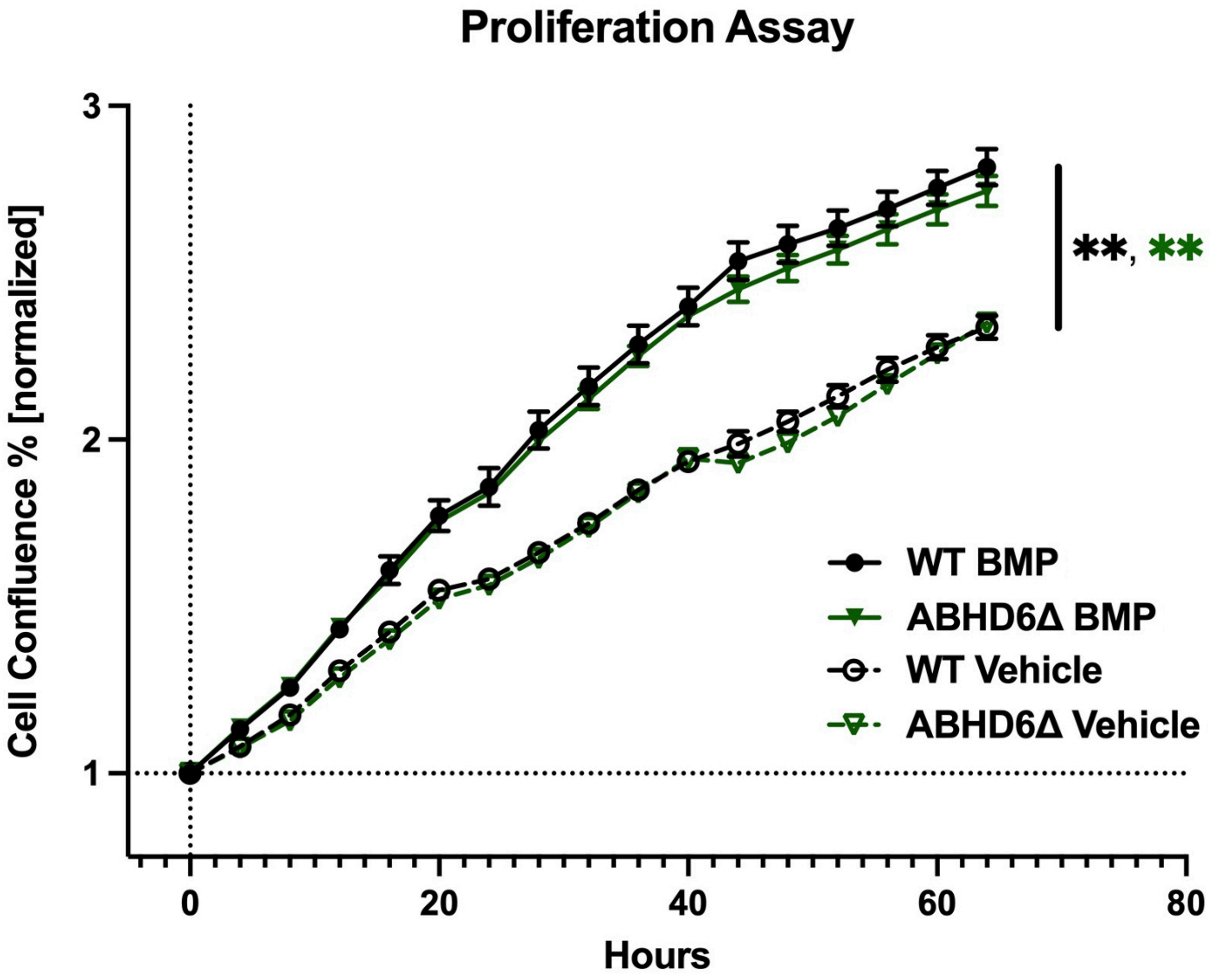


**Supplemental Figure 4. BMP Lipids Promote Proliferation of a Human Hepatoma Cell Line in an ABHD6-Independent Manner.** Huh7 wild type (WT) and ABHD6Δ cells were seeded in 48-well plates and placed in the Incucyte® Live-Cell Analysis System. Cells were cultured in serum free media supplemented with 25 μM bis(monoacylglycerol)phosphates (BMP) versus vehicle (0.1 % DMSO). BMP lipid treatment significantly increased cellular proliferation in both WT and ABHD6Δ cells. No significant differences were seen between the WT and ABHD6Δ cells in the BMP-treatment nor vehicle-treatment groups. Two-way ANOVA, n=4. Experiment performed in biological duplicate (total n=8), and representative graph displayed as mean +/- SEM. *p<0.05, **p<0.01, ***p<0.001

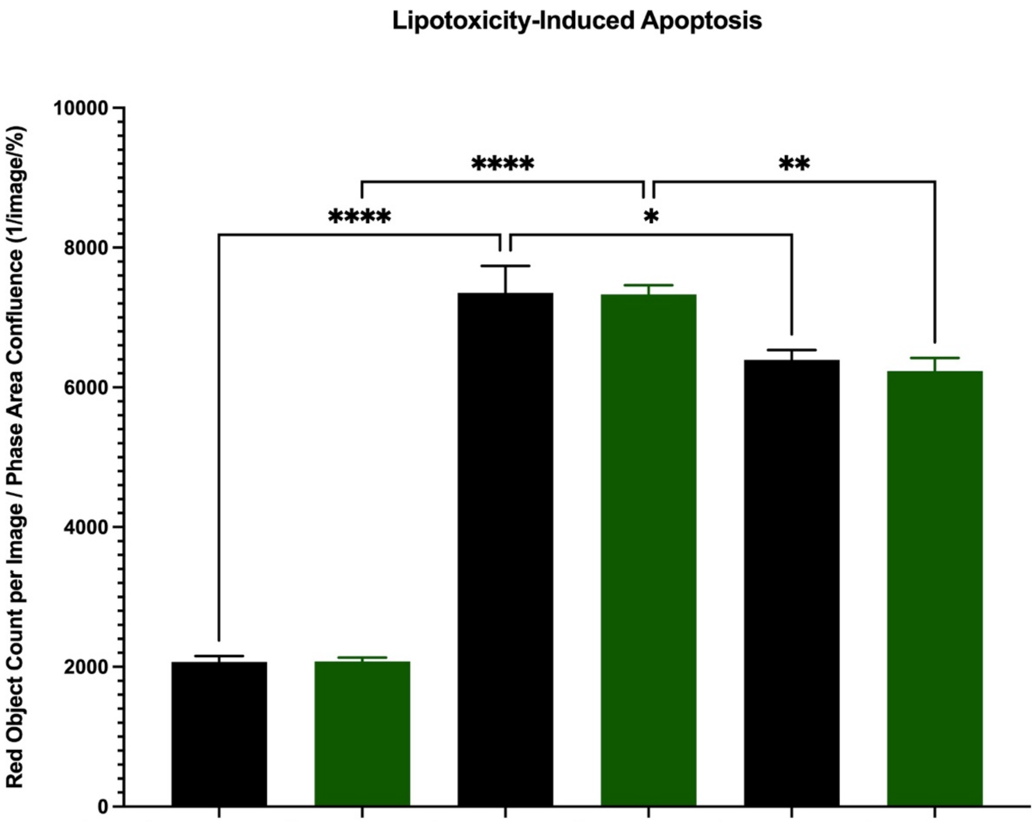


| Genotype | WT | ABHD6Δ | WT | ABHD6Δ | WT | ABHD6Δ |
|---|---|---|---|---|---|---|
| **BMP Treatment** | $Vehicle_{BMP}$ (0.1% DMSO) | $Vehicle_{BMP}$ (0.1% DMSO) | $Vehicle_{BMP}$ (0.1% DMSO) | $Vehicle_{BMP}$ (0.1% DMSO) | 25 μM BMP | 25 μM BMP |
| **PA Treatment** | $Vehicle_{PA}$ (1% IPA + 1% BSA) | $Vehicle_{PA}$ (1% IPA + 1% BSA) | 800 μM PA | 800 μM PA | 800 μM PA | 800 μM PA |

**Supplemental Figure 5. Palmitic Acid-Mediated Lipotoxicity Induces Apoptosis in a Human Hepatoma Cell Line in an ABHD6-Independent Manner.** Huh7 wild type (WT) and ABHD6Δ cells were seeded in 96-well plates and placed in the Incucyte® Live-Cell Analysis System. Cells were treated with 800 μM palmitic acid (PA) versus $vehicle_{PA}$ (1% isopropanol (IPA) + 1% bovine serum albumin (BSA)) for 24 hours. Cells treated with 800 μM PA were supplemented with 25 μM bis(monoacylglycerol)phosphates (BMP) versus $vehicle_{BMP}$ (0.1% DMSO). After 24 hours of treatment, media was replaced and supplemented with a DNA-intercalating dye that binds to the activated caspase-3/7 recognition motif causing cells to fluoresce when caspase-3/7 mediated apoptosis is activated. PA treatment (right-sided four columns) resulted in significantly increased apoptosis in both WT and ABHD6Δ cells compared to $vehicle_{PA}$ alone (left-sided two columns). BMP lipids (right-sided two columns) significantly decreased apoptosis in both WT and ABHD6Δ cells treated with palmitic acid compared to $vehicle_{BMP}$ alone (middle two columns). No significant differences were seen between the WT and ABHD6Δ cells in all groups. One-way ANOVA, n=5. Experiment performed in biological duplicate (total n=10), results combined and resultant graph displayed as mean +/- SEM. *$p<0.05$, **$p<0.01$, ***$p<0.001$, ****$p<0.0001$.

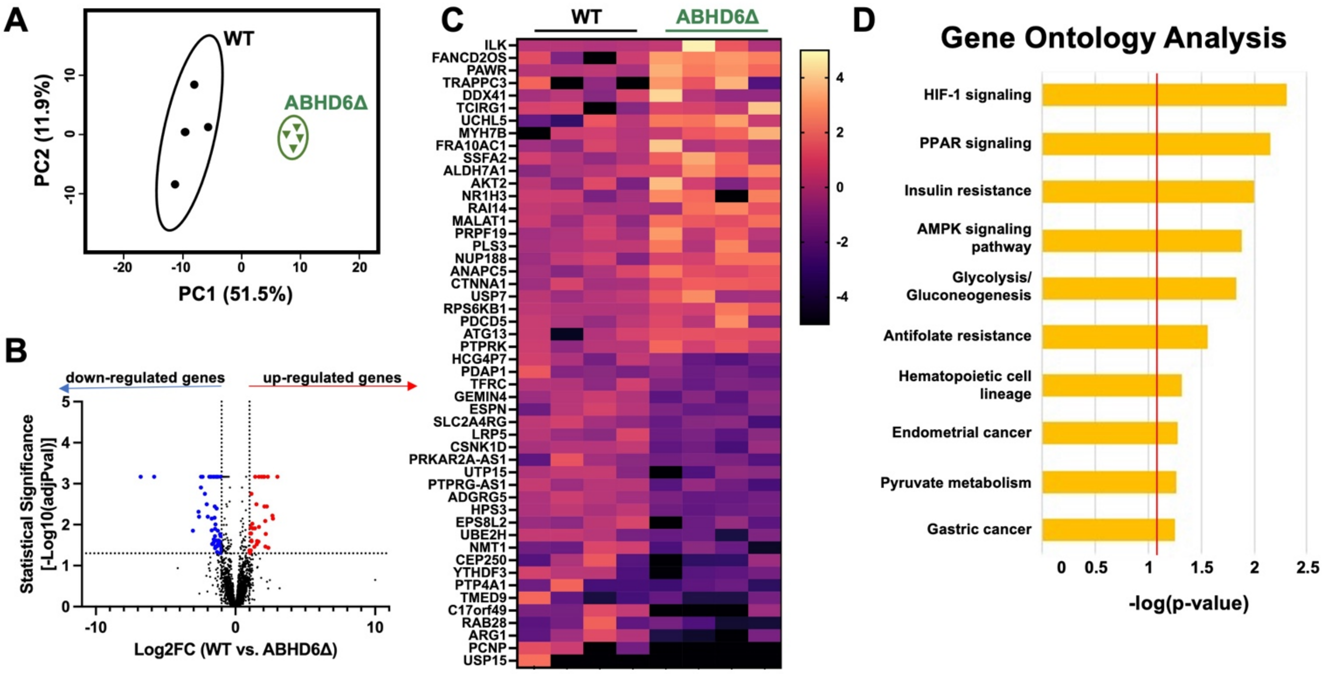


**Supplemental Figure 6. ABHD6 Knockout Alters Lipid Metabolism Pathways in a Human Hepatoma Cell Line.** Huh7 wild type (WT) and ABHD6Δ cells were cultured in basal media for 24 hours and harvested for RNAseq analysis. **(A)** Principal component analysis showed distinct clustering of WT and ABHD6Δ cells. **(B)** Volcano plot of gene expression changes (n = 4; genes with an absolute value log2 fold change greater than 1 and adjusted p-value < 0.05 were considered significantly differentially expressed), **(C)** and heat map of row-normalized expression for the top 50 differentially expressed genes demonstrate strong gene expression changes between groups. **(D)** Gene ontology analysis was notable for differences involving lipid signaling, with changes observed in HIF-1, PPAR, and AMPK signaling pathways.

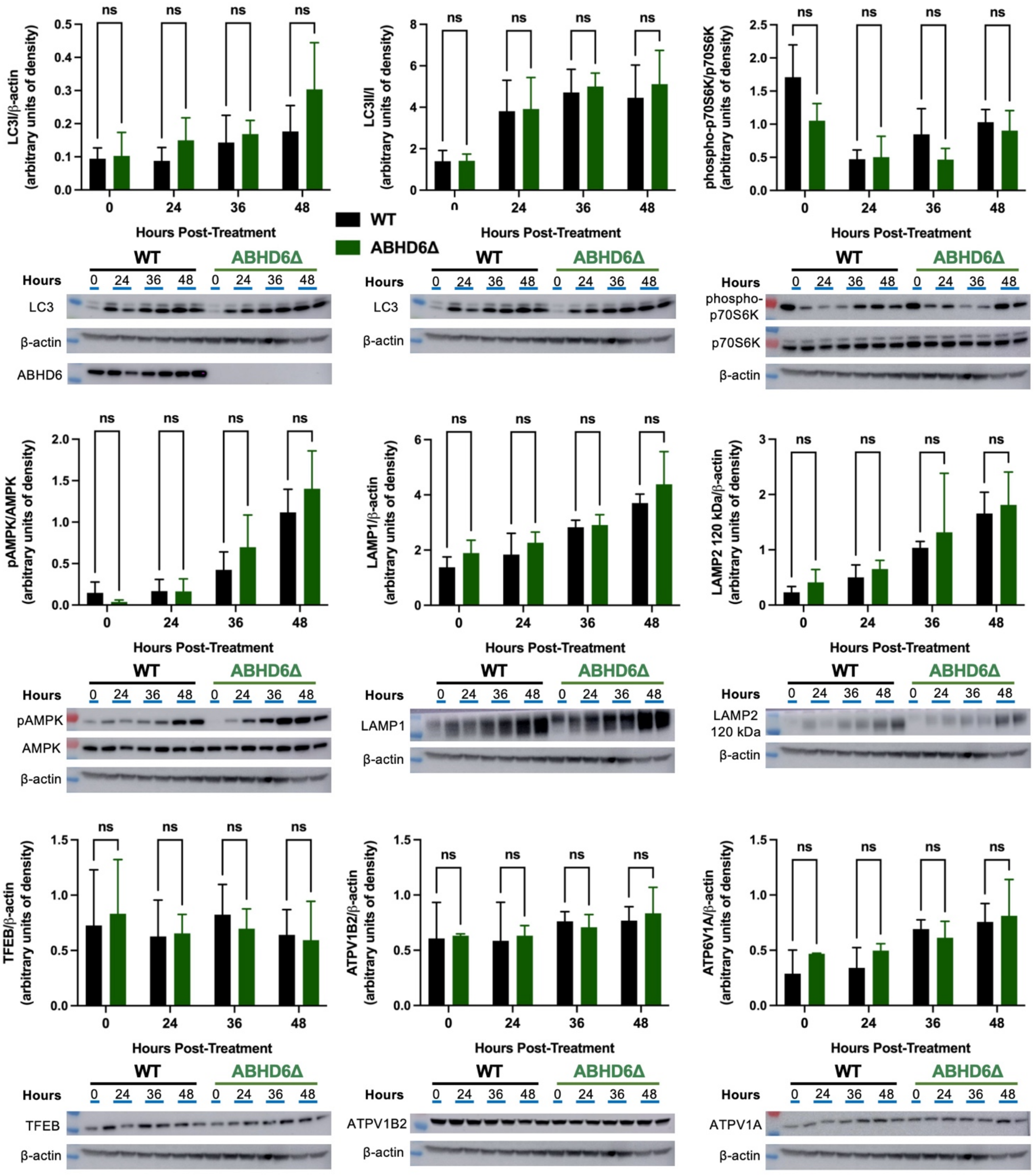


**Supplemental Figure 7. Lysosomal Pathway and Autophagy-Relevant Protein Expression in a Human Hepatoma Cell Line Subjected to Palmitic Acid-Mediated Lipotoxicity.** Huh7 wild type (WT) and ABHD6Δ cells were treated with 800 μM palmitic acid (PA) for up to 48 hours. Cells were harvested for western blot analysis. Densitometry analysis (above) and representative western blots (below) are shown. Multiple Mann-Whitney tests, n=2. Experiment performed in biological duplicate (total n=6), results combined and resultant graph displayed as mean +/- SD. *p<0.05, **p<0.01, ***p<0.001.

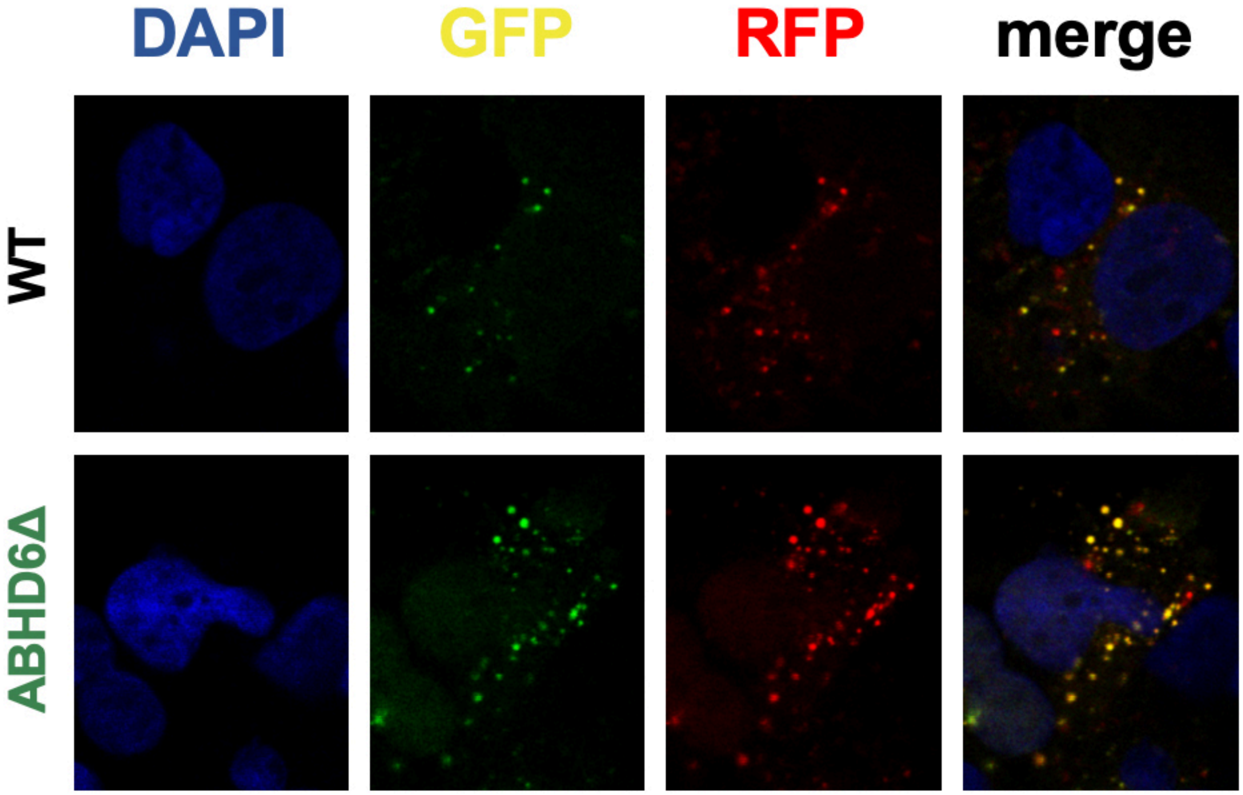


**Supplemental Figure 8. Autophagic Flux Measured Using an RFP-GFP-LC3 Reporter in a Human Hepatoma Cell Line Subjected to Palmitic Acid-Mediated Lipotoxicity.** Huh7 wild type (WT) and ABHD6Δ cells were treated with 800 μM palmitic acid (PA) for 24 hours, and the RFP-GFP-LC3 reporter was used to study autophagic flux. Representative confocal microscopy images are shown (from left to right: DAPI, GFP, RFP, merge).

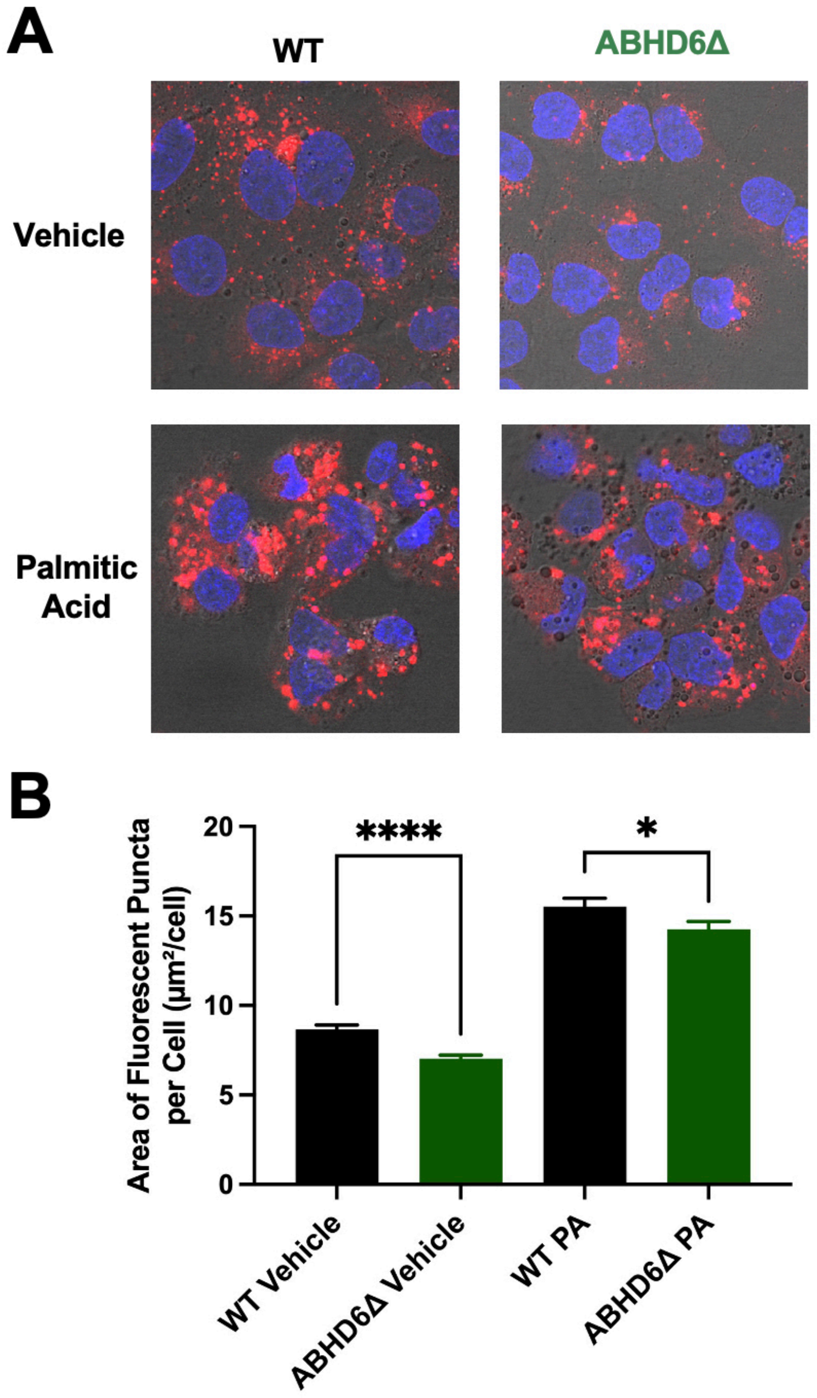


**Supplemental Figure 9. LysoTracker Staining in a Human Hepatoma Cell Line Subjected to Palmitic Acid-Mediated Lipotoxicity.** Huh7 wild type (WT) and ABHD6Δ cells were treated with 800 μM palmitic acid (PA) versus vehicle (1% isopropanol (IPA) + 1% bovine serum albumin (BSA)) alone for 24 hours. LysoTracker Deep Red staining was performed. **(A)** Representative images and **(B)** quantification demonstrate an ABHD6-independent increase in LysoTracker staining with PA treatment. Decreased LysoTracker staining was seen in ABHD6Δ cells treated with both PA treatment and vehicle alone. Unpaired t-test, n=2. Experiment performed in biological duplicate, and results combined and resultant graph displayed as mean +/- SEM. $^*p<0.05$, $^{**}p<0.01$, $^{***}p<0.001$.

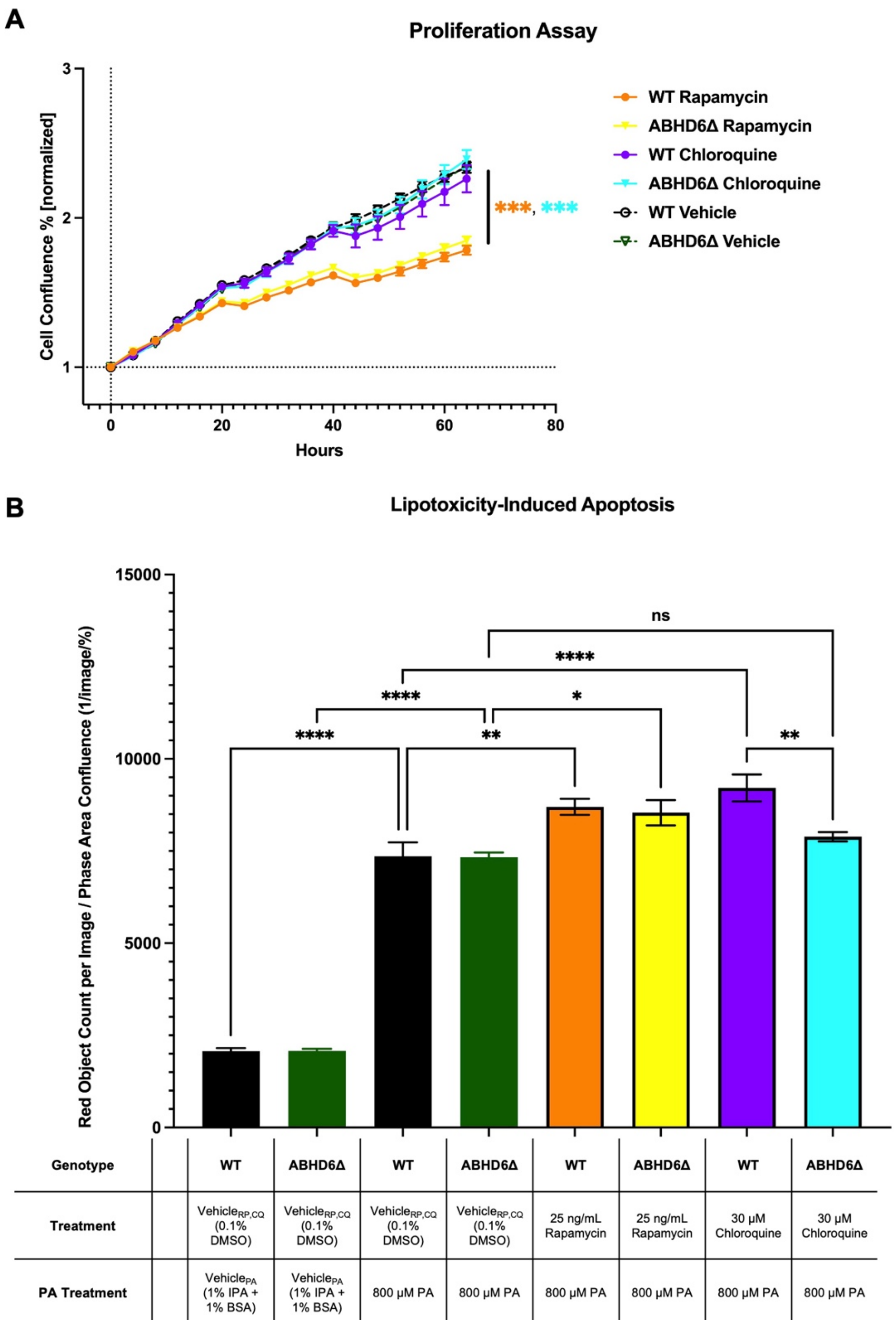


**Supplemental Figure 10. Effects of Rapamycin and Chloroquine on Cell Proliferation and Palmitic Acid-Mediated Lipotoxicity-Induced Apoptosis of a Human Hepatoma Cell Line.** **(A)** Huh7 wild type (WT) and ABHD6Δ cells were seeded in 48-well plates and placed in the Incucyte® Live-Cell Analysis System. Cells were cultured in serum free media supplemented with 25 ng/ml rapamycin, 30 µM chloroquine, or vehicle (0.1 % DMSO) alone. Rapamycin treatment significantly reduced cell proliferation in both WT and ABHD6Δ cells. No significant differences were seen between the WT and ABHD6Δ cells in all groups. Two-way ANOVA, n=4. Experiment performed in biological duplicate (total n=8), and representative graph displayed. **(B)** Huh7 WT and ABHD6Δ cells were seeded in 96-well plates and placed in the Incucyte® Live-Cell Analysis System. Cells were treated with 800 µM palmitic acid (PA) versus vehicle$_{PA}$

(1% isopropanol (IPA) + 1% bovine serum albumin (BSA)) for 24 hours. Cells treated with 800 μM PA were supplemented with 25 ng/ml rapamycin (RP), 30 μM chloroquine (CQ), or $vehicle_{RP,CQ}$ (0.1 % DMSO) alone. After 24 hours of treatment, media was replaced and supplemented with a DNA-intercalating dye designed to fluoresce upon cleavage by activated caspase-3/7. PA treatment (right-sided six columns) resulted in significantly increased apoptosis in both WT and ABHD6Δ cells compared to $vehicle_{PA}$ alone (left-sided two columns). A significant increase in apoptosis was seen with rapamycin treatment compared to treatment with $vehicle_{RP,CQ}$ alone in an ABHD6-independent manner. Chloroquine treatment significantly increased apoptosis in WT cells but not ABHD6Δ cells. No significant differences were seen between the WT and ABHD6Δ cells in all groups except with chloroquine treatment. One-way ANOVA, n=5. Experiment performed in biological duplicate (total n=10), results combined and resultant graph displayed. Graphs displayed as mean +/- SEM. *p<0.05, **p<0.01, ***p<0.001, ****p<0.0001.
